## Supplementary Information for "A Systematic Evaluation of Single Cell RNA-Seq Analysis Pipelines"

by

Vieth et al.

### Supplementary Figures

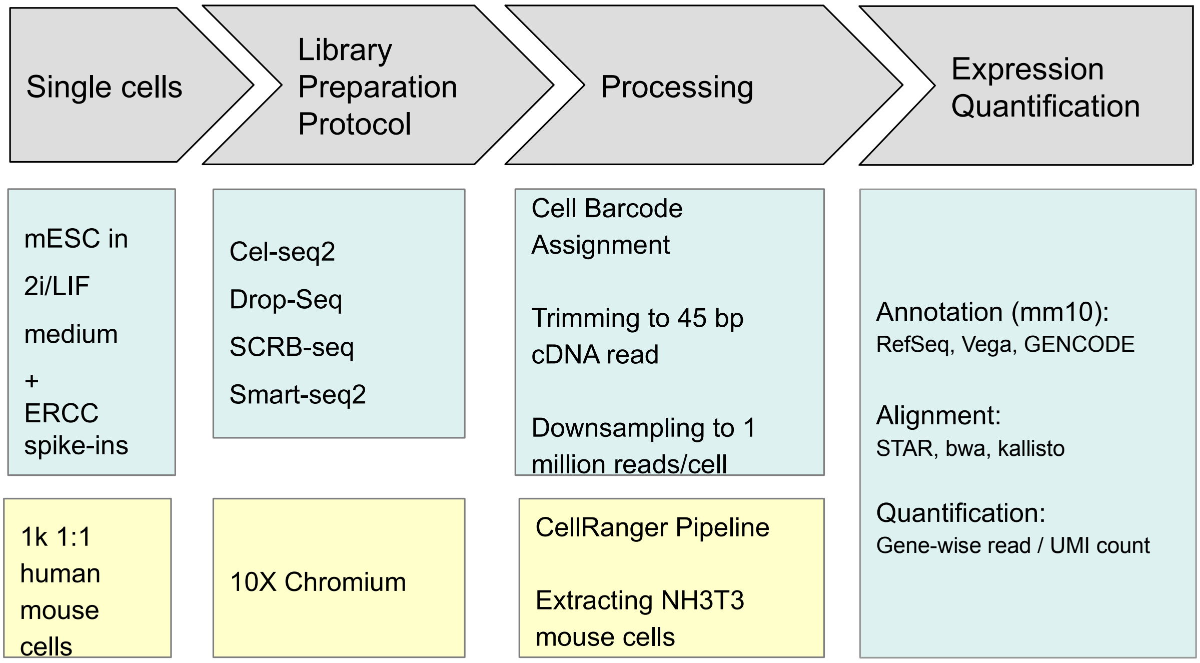


**Supplementary Figure 1:** Schematic overview of scRNA-seq data sets. In our comparison, we included the gene expression profiles of mouse embryonic stem cells (mESC) as published in[1]. Briefly, the mESC were cultured under two inhibitor/leukemia inhibitory factor (2i /LIF) conditions to ensure rather homogeneous cell populations[2]. We selected four scRNA-seq methods (Smart-seq2, SCRB-seq, Drop-seq, CEL-seq2) that were used to construct libraries in two independent replicate batches. In addition, 92 poly-adenylated synthetic RNA transcripts of known concentration designed by the External RNA Control Consortium (ERCCs)[3] were spiked in for all methods except Drop-seq. All raw sequencing reads were cut and downsampled to 45 base long one million cDNA reads per cell. We included two commonly used expression quantification approaches, namely reference-guided alignment in STAR and bwa as well as pseudoalignment in kallisto in combination with three annotations. Furthermore, we downloaded a scRNA-seq data set from 10X Genomics Support, namely the 1k 1:1 Mixture of Fresh Frozen Human (HEK293T) and Mouse (NH3T3) Cells[4] generated using the v2 gene expression chemistry. We proceeded with approx. 400 mouse cells with 70000 reads/cell on average.

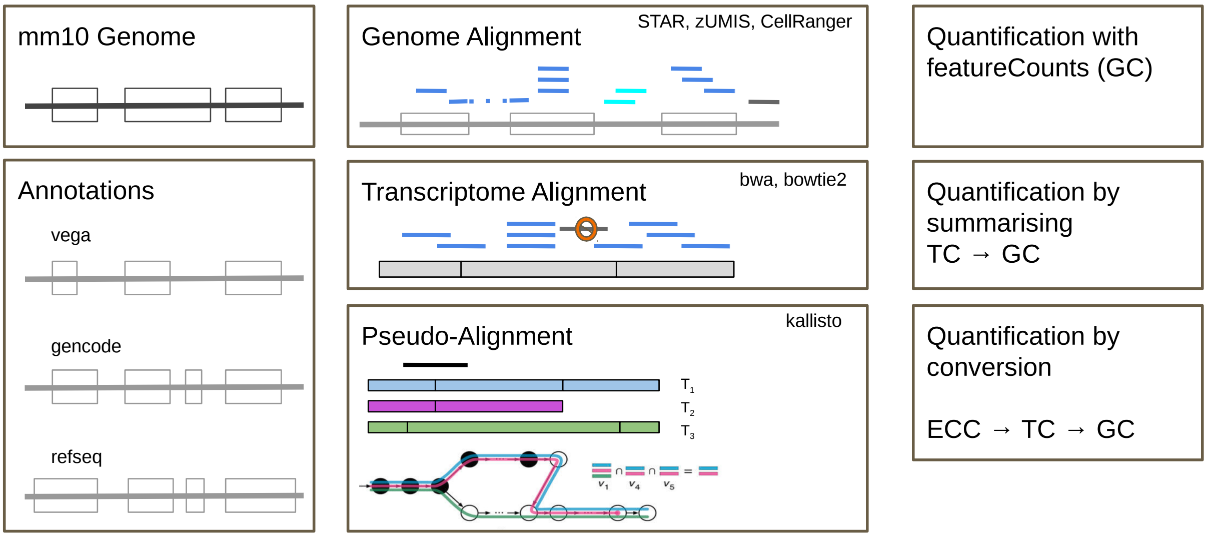


**Supplementary Figure 2:** Schematic overview of alignment and annotation. **Left** The annotation RefSeq, Vega and GENCODE for the mm10 genome **Middle** Alignment of reads to the genome using STAR allowing alignment to genic and intergenic sequences; Alignment of reads to the transcriptome using BWA; Pseudo-alignment of reads to de Bruijn Graph representation of the transcriptome using kallisto (figure adapted from[5]). **Right** STAR: Estimation of expression as read/UMI counts per gene (GC) using featureCounts for genome alignments; BWA: Estimation of expression as read/UMI counts per gene by summarising counts over transcripts (TC) belonging to one gene; kallisto: Estimation of expression given as equivalence class counts (ECC) converted to trancript counts (TC) and summarised to gene counts (GC).

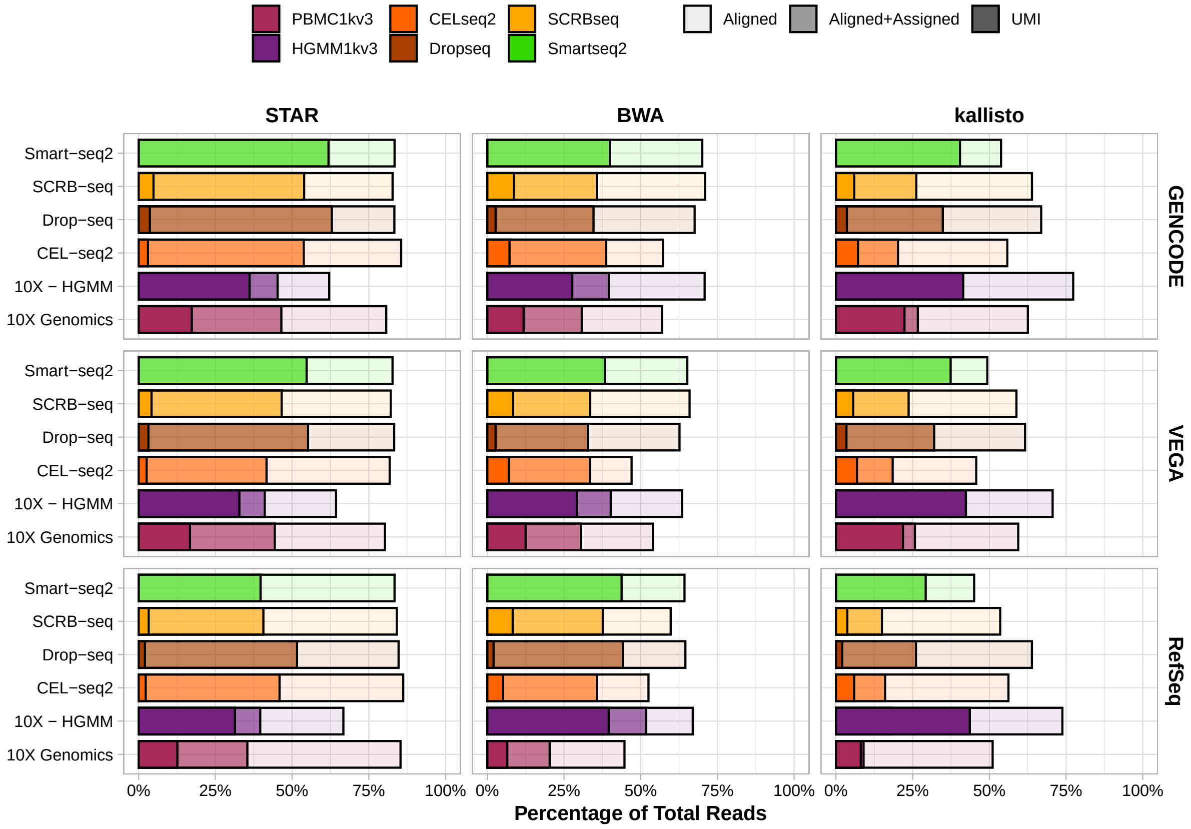


**Supplementary Figure 3:** Read Alignment and Assignment Rates per Library Preparation Protocol stratified over Aligner and Annotation. The lighter shade represents the percentage of the total reads that could be aligned and the darker shade the percentage that also was uniquely assigned.

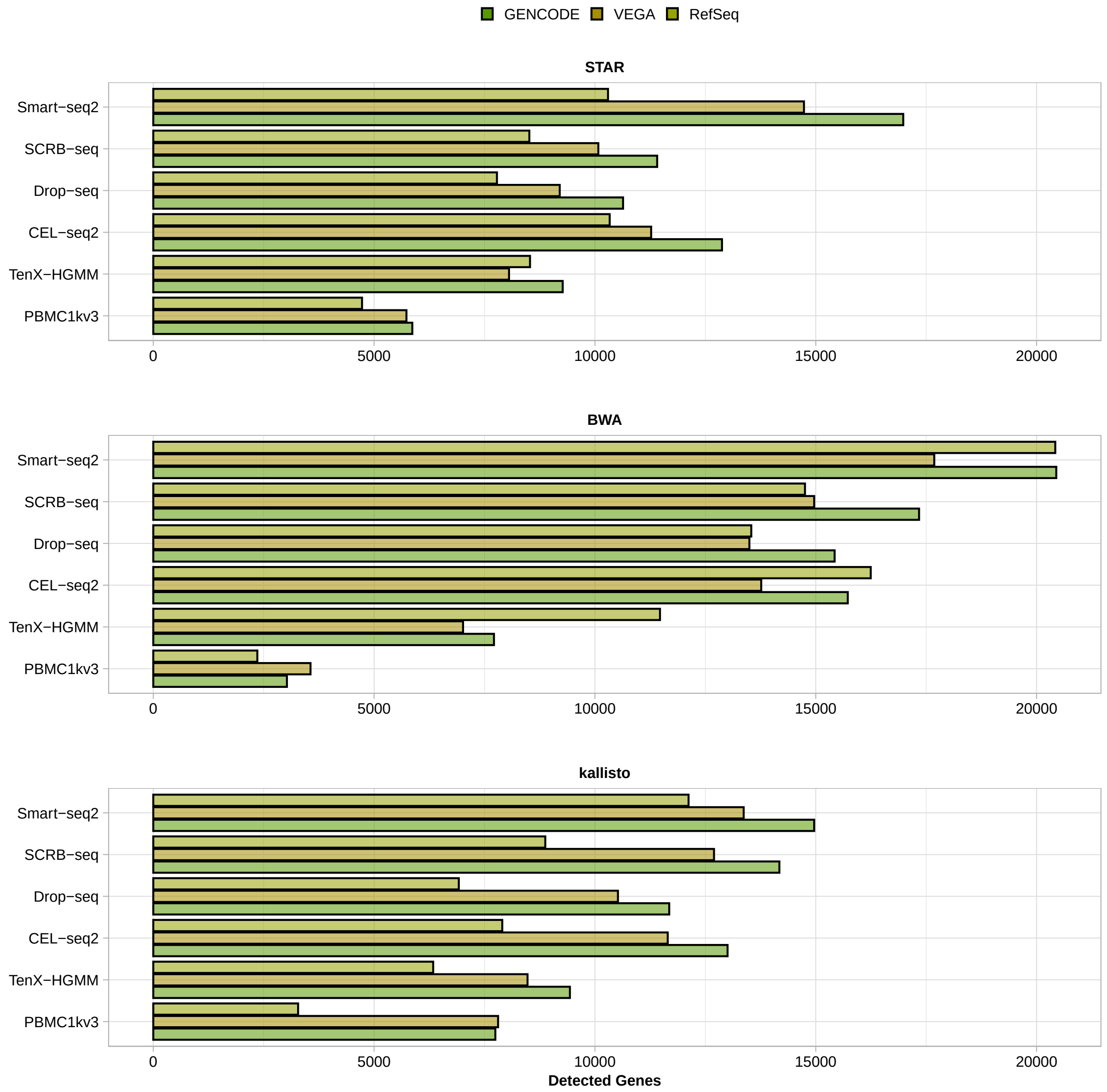


**Supplementary Figure 4:** Gene Detection Rates. Number of genes detected per Library Preparation Protocol stratified over Aligner and Annotation (i.e. at least 10% nonzero expression values).

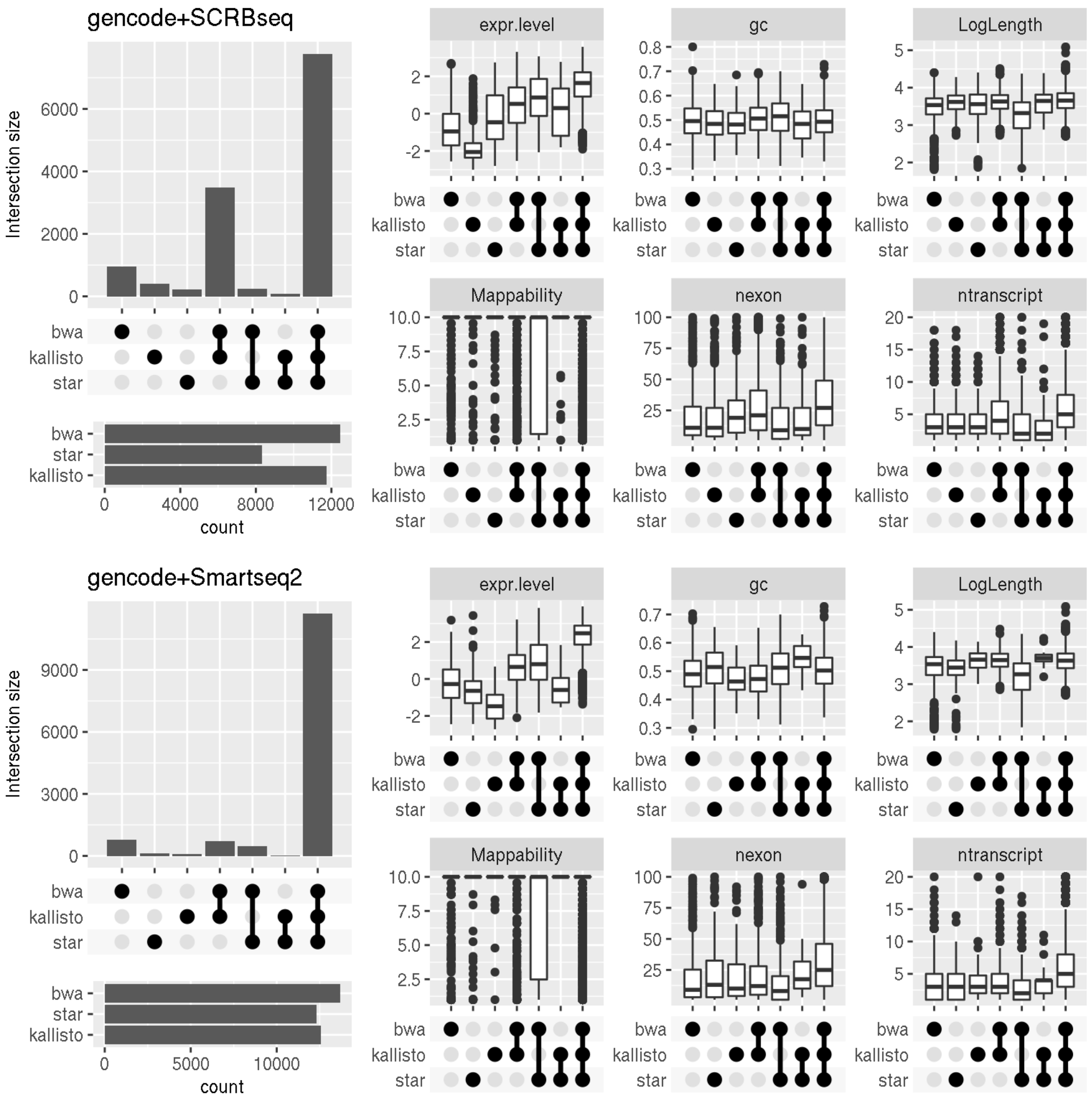


**Supplementary Figure 5:** Properties of genes detected by the different mapping strategies. Mappability is represented as $10^{M25}$, where M25 is the lower quartile of the mappability scores[6] across the gene. All other gene properties were extracted from the corresponding annotation file. Genes detected by all three mappers tend to have higher expression levels. The only other consistent pattern is that genes only detected by kallisto have on average a lower expression and genes that escape detection by kallisto have slightly lower mappability. Box and whisker plot with centre line = median, bounds of box = 25th and 75th percentile, whiskers = 1.5 * interquartile range from the lower and upper bounds of the box.

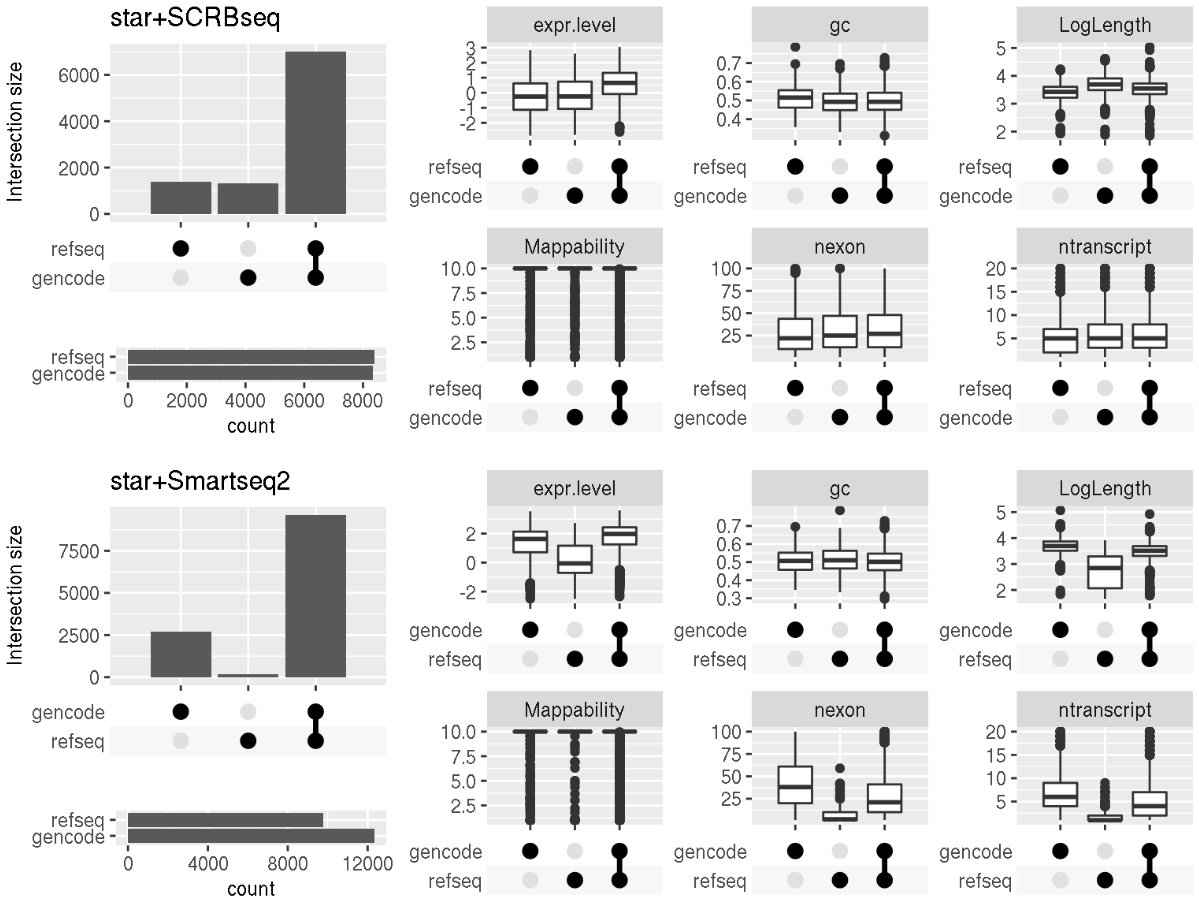


**Supplementary Figure 6:** Differences in genes detected with RefSeq and Gencode annotation. We used the matching between RefSeq and Gencode transcript annotations that is provided by Gencode and then summarise detection at the gene level. Comparing the genes found using Gencode vs. RefSeq annotation, we find that for the 3-prime method SCRB-seq both annotations yield approximately the same number of genes, but ∼1,000 appear annotation-specific. The Gencode annotation is more comprehensive, in that it contains more and often longer transcripts and indeed the Gencode-specific genes are longer and thus 3 mapping reads are easily lost. The RefSeq specific transcripts are more puzzling and the only distinguishing feature that we see is that they appear more GC-rich. For full-length data generated with Smart-seq2, Gencode detects 2,500 more genes than RefSeq and there are almost no RefSeq specific genes. Genes detected with Gencode only are longer and have on average more exons and transcripts. Box and whisker plot with centre line = median, bounds of box = 25th and 75th percentile, whiskers = 1.5 * interquartile range from the lower and upper bounds of the box.

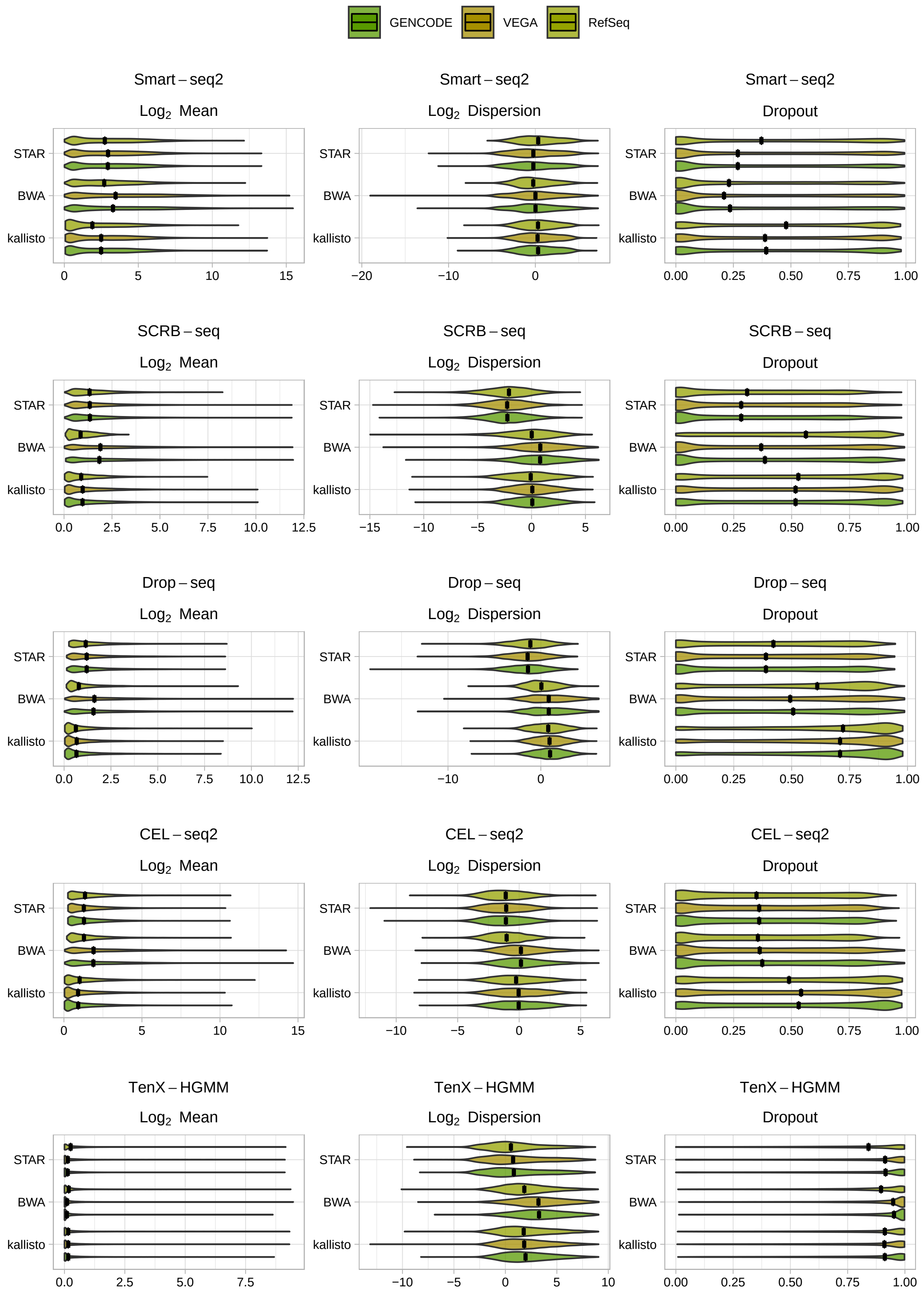


**Supplementary Figure 7:** Distribution Estimates. Estimated mean expression, dispersion and dropout rates per Library Preparation Protocol stratified over Aligner and Annotation. Black line indicates median value.

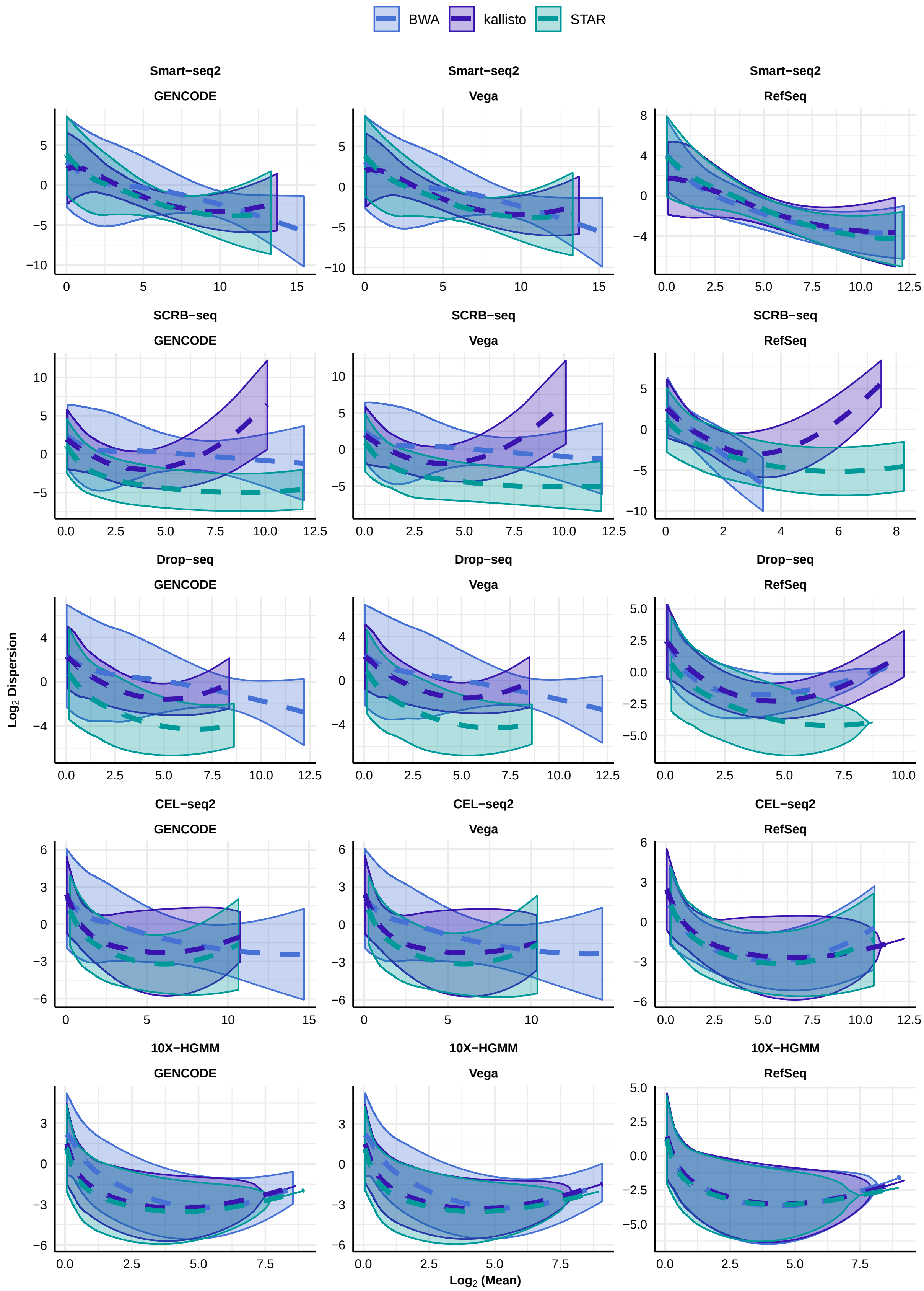


**Supplementary Figure 8:** Distributional Fittings for simulations. Mean-Dispersion Fitting Line applying a cubic smoothing spline with 95% variability bands per Library Preparation Protocol stratified over Aligner and Annotation.

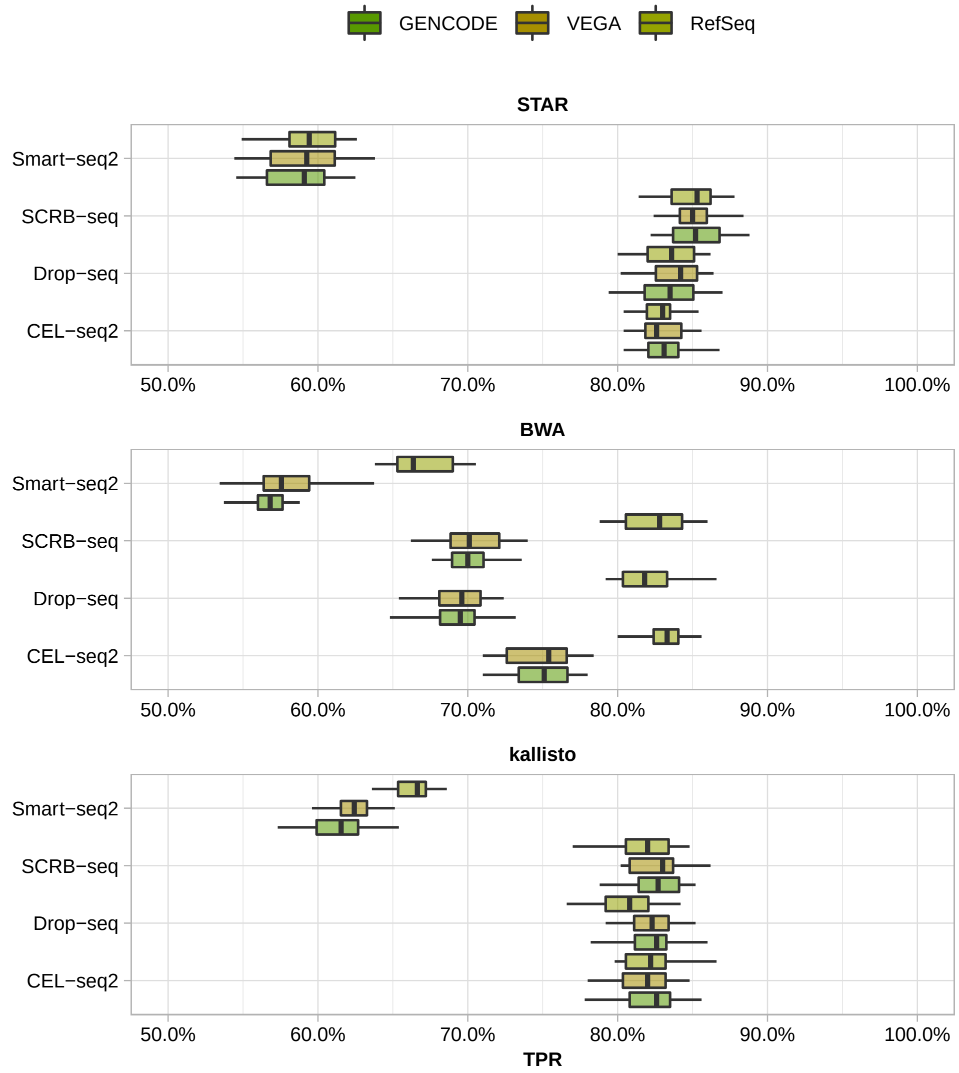


**Supplementary Figure 9:** The effect of quantification choices on detecting differential expression. The expression of 10,000 genes over 768 cells (384 cells per group) was simulated and 5% of the genes were differentially expressed following an asymmetric narrow gamma distribution. Any gene correctly called differentially expressed at FDR 10% contributed to the True Positive Rate (TPR). The TPR per library preparation method stratified over aligner and annotation is plotted. Box and whisker plot with centre line = median, bounds of box = 25th and 75th percentile, whiskers = 1.5 * interquartile range from the lower and upper bounds of the box.

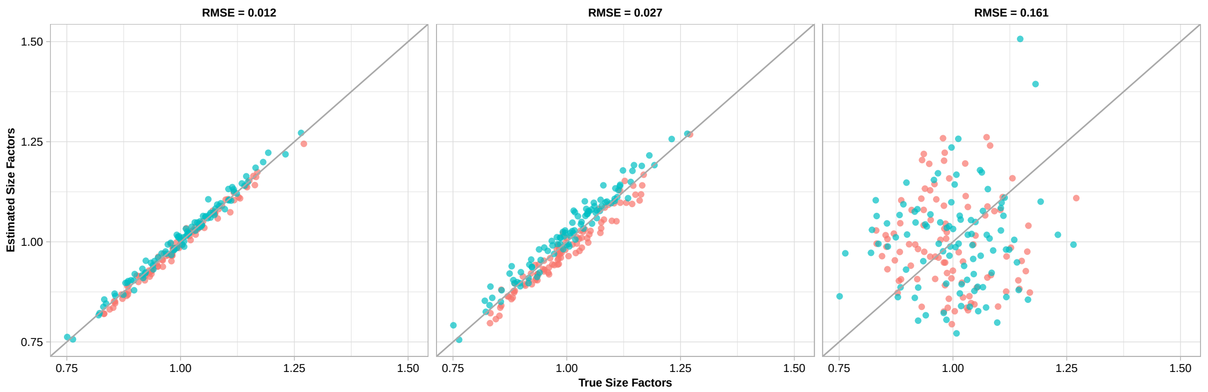


**Supplementary Figure 10:** Illustration for RMSE evaluation of library size factors. The expression of 10,000 genes over 768 cells (384 cells per group (red and cyan points) were simulated and 20% of the genes were differentially expressed following an asymmetric narrow gamma distribution. To compare the estimated library size factors with the simulated library size factors, the factors were centred and scaled. The root mean squared error (RMSE) of a robust linear regression represents the deviation between estimated and simulated size factors.

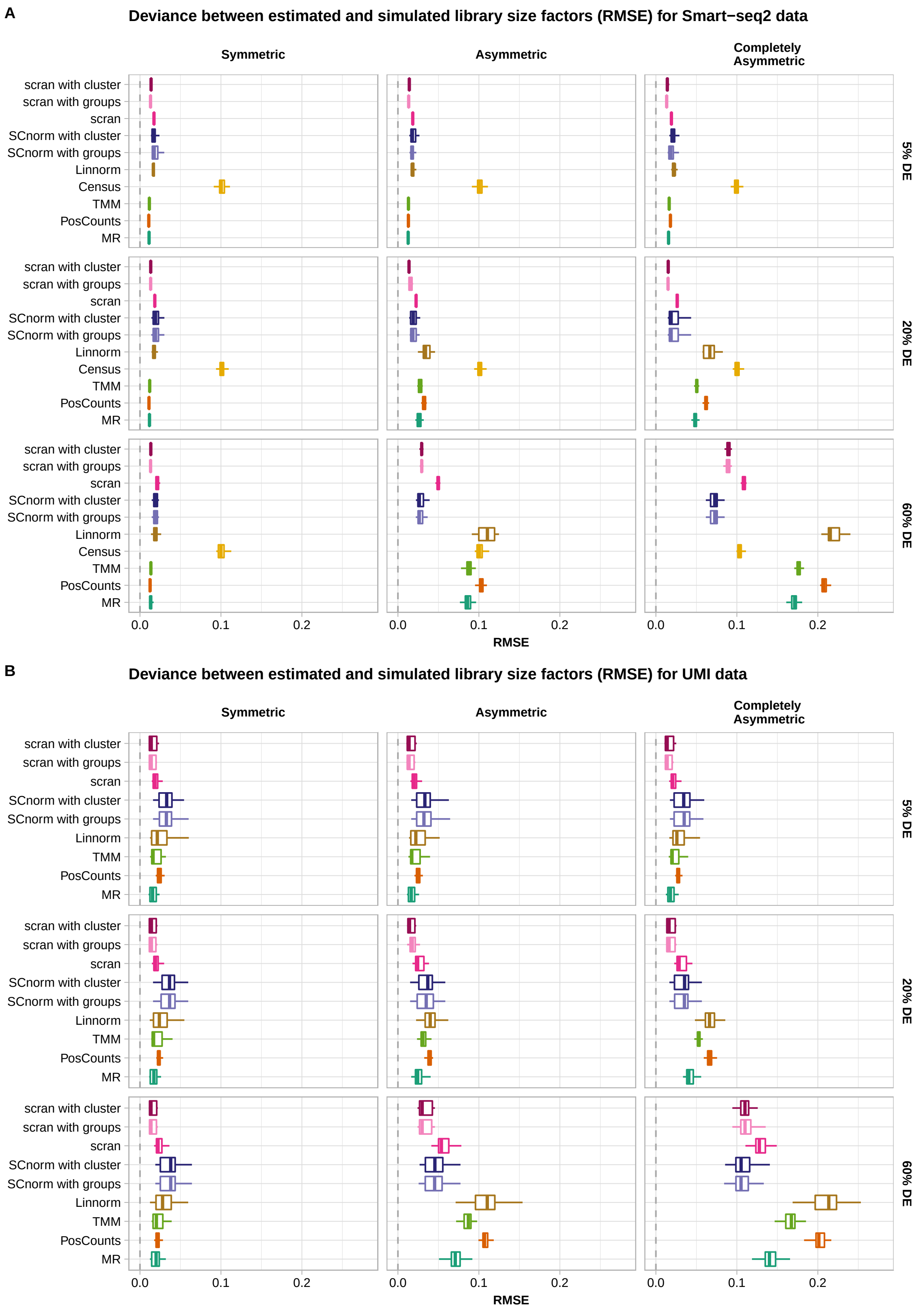


**Supplementary Figure 11:** Deviation between simulated and estimated library size factors. The expression of 10,000 genes over 768 cells (384 cells per group) were simulated and 5%, 20% or 60% of the genes were differentially expressed following a symmetric, an asymmetric or a completely asymmetric narrow gamma distribution. The root mean squared error (RMSE) of estimated scaling factors per normalisation method is plotted.Box and whisker plot with centre line = median, bounds of box = 25th and 75th percentile, whiskers = 1.5 * interquartile range from the lower and upper bounds of the box. **A** Smart-seq2 data **B** UMI data.

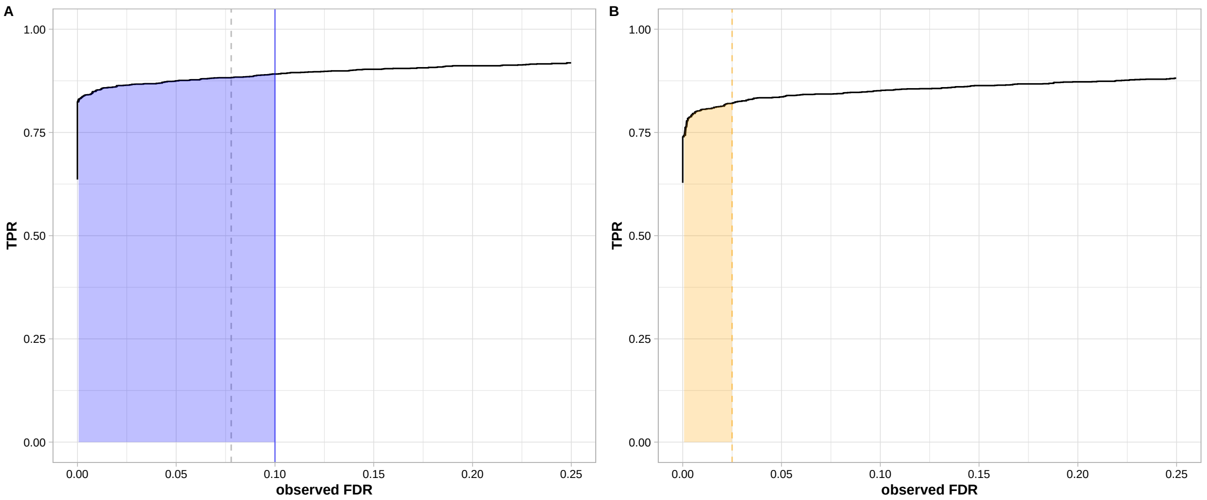


C

| TP | FP | TN | FN | TPR | observed FDR | threshold FDR |
|---|---|---|---|---|---|---|
| 1687 | 12 | 7988 | 313 | 0.84 | 0.0071 | 0.01 |
| 1716 | 23 | 7977 | 284 | 0.86 | 0.0132 | 0.02 |
| 1729 | 45 | 7955 | 271 | 0.86 | 0.0254 | 0.03 |
| 1735 | 60 | 7940 | 265 | 0.87 | 0.0334 | 0.04 |
| 1739 | 77 | 7923 | 261 | 0.87 | 0.0424 | 0.05 |
| 1747 | 83 | 7917 | 253 | 0.87 | 0.0454 | 0.06 |
| 1752 | 98 | 7902 | 248 | 0.88 | 0.053 | 0.07 |
| 1755 | 114 | 7886 | 245 | 0.88 | 0.061 | 0.08 |
| 1762 | 126 | 7874 | 238 | 0.88 | 0.0667 | 0.09 |
| 1765 | 149 | 7851 | 235 | 0.88 | 0.0778 | 0.1 |
| 1771 | 170 | 7830 | 229 | 0.89 | 0.0876 | 0.11 |
| 1779 | 190 | 7810 | 221 | 0.89 | 0.0965 | 0.12 |
| 1783 | 199 | 7801 | 217 | 0.89 | 0.1004 | 0.13 |
| 1788 | 214 | 7786 | 212 | 0.89 | 0.1069 | 0.14 |
| 1790 | 226 | 7774 | 210 | 0.9 | 0.1121 | 0.15 |

D

| TP | FP | TN | FN | TPR | observed FDR | threshold FDR |
|---|---|---|---|---|---|---|
| 1678 | 93 | 7907 | 322 | 0.84 | 0.0525 | 0.01 |
| 1688 | 148 | 7852 | 312 | 0.84 | 0.0806 | 0.02 |
| 1705 | 198 | 7802 | 295 | 0.85 | 0.104 | 0.03 |
| 1712 | 246 | 7754 | 288 | 0.86 | 0.1256 | 0.04 |
| 1726 | 296 | 7704 | 274 | 0.86 | 0.1464 | 0.05 |
| 1729 | 332 | 7668 | 271 | 0.86 | 0.1611 | 0.06 |
| 1735 | 371 | 7629 | 265 | 0.87 | 0.1762 | 0.07 |
| 1742 | 404 | 7596 | 258 | 0.87 | 0.1883 | 0.08 |
| 1745 | 437 | 7563 | 255 | 0.87 | 0.2003 | 0.09 |
| 1748 | 473 | 7527 | 252 | 0.87 | 0.213 | 0.1 |
| 1756 | 522 | 7478 | 244 | 0.88 | 0.2291 | 0.11 |
| 1758 | 562 | 7438 | 242 | 0.88 | 0.2422 | 0.12 |
| 1764 | 597 | 7403 | 236 | 0.88 | 0.2529 | 0.13 |
| 1767 | 636 | 7364 | 233 | 0.88 | 0.2647 | 0.14 |
| 1773 | 662 | 7338 | 227 | 0.89 | 0.2719 | 0.15 |

**Supplementary Figure 12:** Illustration for pAUC calculation used as an evaluation metric for performance. The expression of 10,000 genes estimated from SCRB-seq data over 768 cells (384 cells per group were simulated and 20% of the genes were differentially expressed following an asymmetric narrow gamma distribution. The power to detect DE (TPR) versus observed FDR based on DE-testing with limma-trend using scran with clustering (**A**) or Median-Ratio (**B**) is plotted[7]. The dashed line indicates the level at the which the observed stays below the nominal level of 10% which defines the right side boundary for the partial area under this curve[8]. The corresponding proportion and rates for a selection of nominal FDR thresholds are given in **C, D**. Since we do not want to punish conservative FDR control of methods, the test results using scran normalisation for example is extended to observed FDR of 10% whereas median ratio only ensures FDR control at a lower observed FDR.

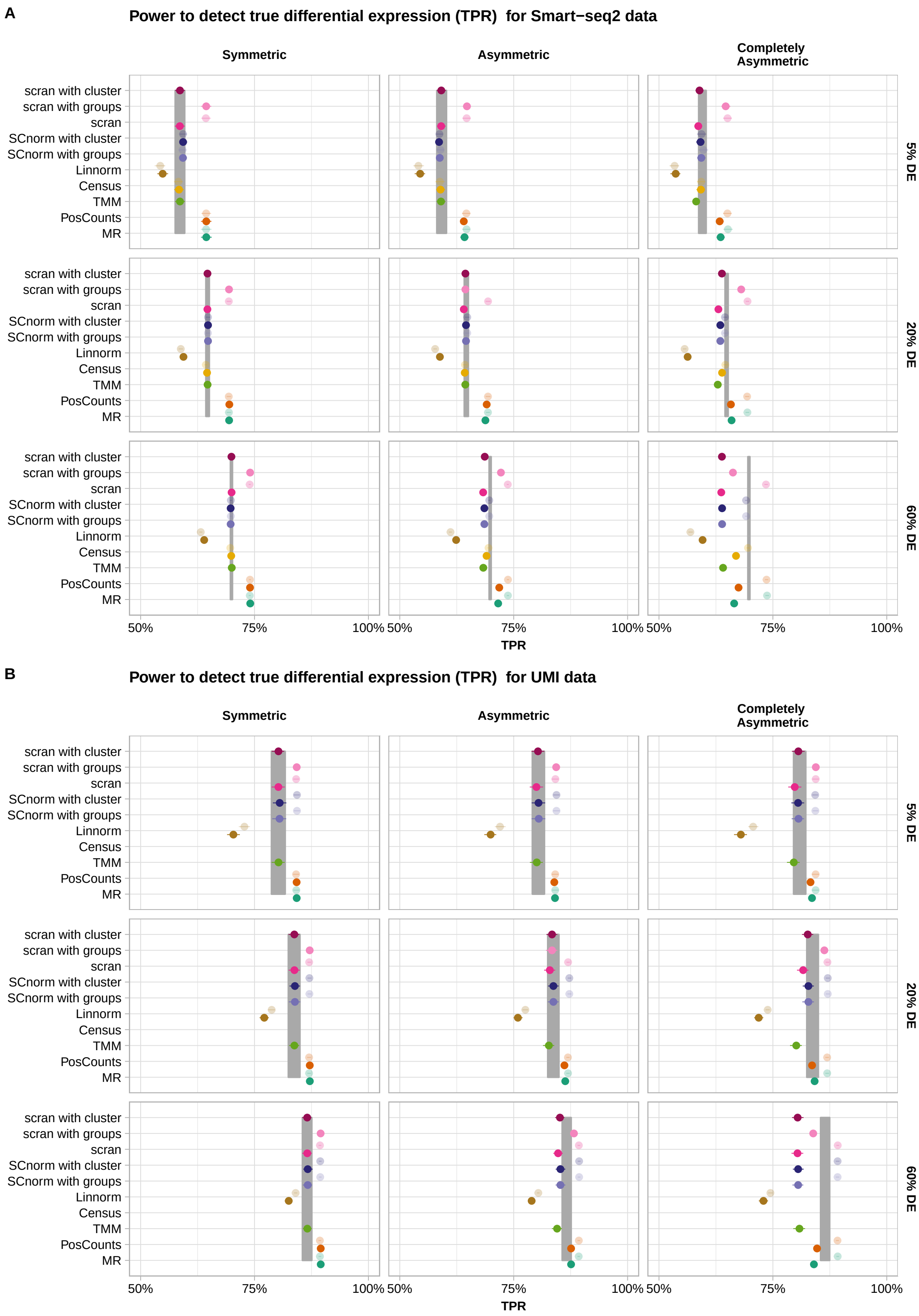


**Supplementary Figure 13:** Power to detect true differential expression per normalisation method. The expression of 10,000 genes over 768 cells (384 cells per group) were simulated and 5%, 20% or 60% of the genes were differentially expressed following a symmetric, an asymmetric or a completely asymmetric narrow gamma distribution. The power to detect DE (mean TPR ± s.d.) based on DE-testing with limma-trend per normalisation method is plotted. The lighter shade indicates the usage of spike-ins for normalisation. The grey ribbon indicates the TPR given simulated size factors (mean TPR ± s.d.). **A** Smart-seq2 data **B** UMI data.

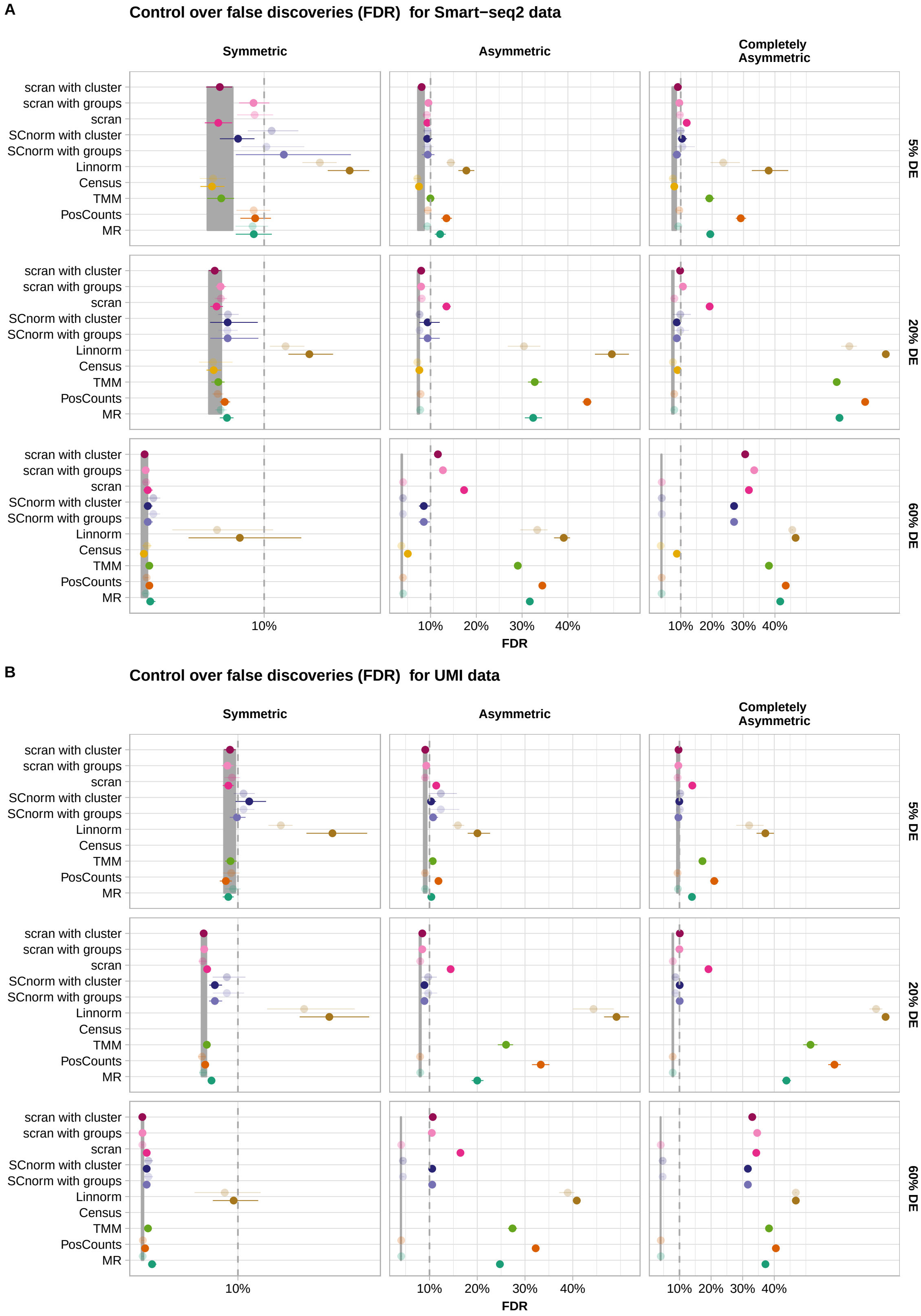


**Supplementary Figure 14:** Control over false discoveries per normalisation method. The expression of 10,000 genes over 768 cells (384 cells per group) were simulated and 5%, 20% or 60% of the genes were differentially expressed following a symmetric, an asymmetric or a completely asymmetric narrow gamma distribution. The BH-FDR control based on DE-testing with limma-trend per normalisation method is plotted (mean FDR ± s.d.). The lighter shade indicates the usage of spike-ins for normalisation. The dashed line indicates the nominal FDR level of 10%. The grey ribbon indicates the FDR given simulated size factors (mean FDR ± s.d.).
**A** Smart-seq2 data **B** UMI data.

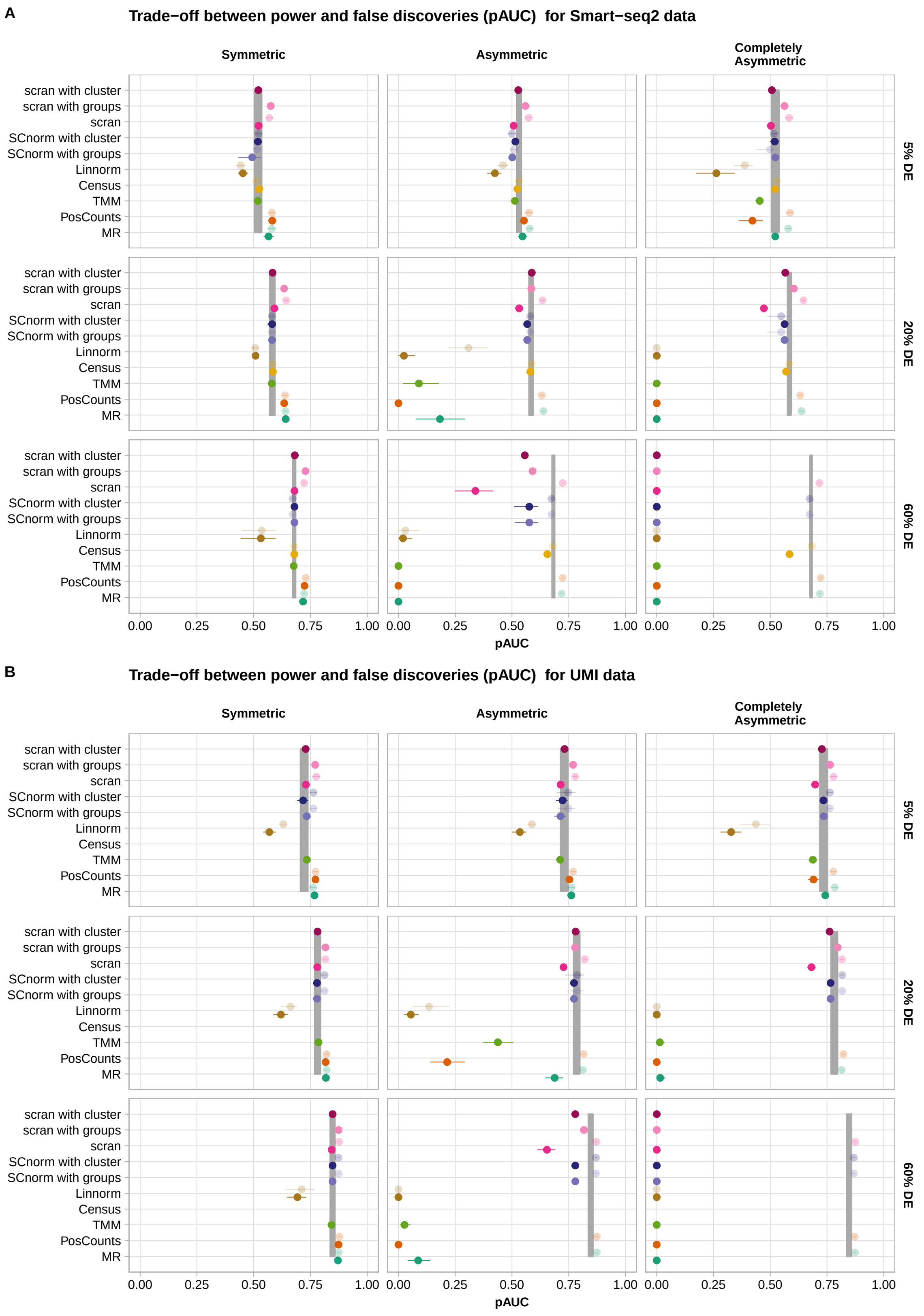


**Supplementary Figure 15:** Trade-off between power and false discoveries per normalisation method. The expression of 10,000 genes over 768 cells (384 cells per group) were simulated and 5%, 20% or 60% of the genes were differentially expressed following a symmetric, an asymmetric or a completely asymmetric narrow gamma distribution. The discriminatory ability determined by the partial area under the curve (pAUC) based on DE-testing with limma-trend for normalisation per DE-setup is plotted (mean pAUC ± s.d.). The lighter shade indicates the usage of spike-ins for normalisation. The grey ribbon indicates the pAUC given simulated size factors (mean pAUC ± s.d.).
**A** Smart-seq2 data **B** UMI data.

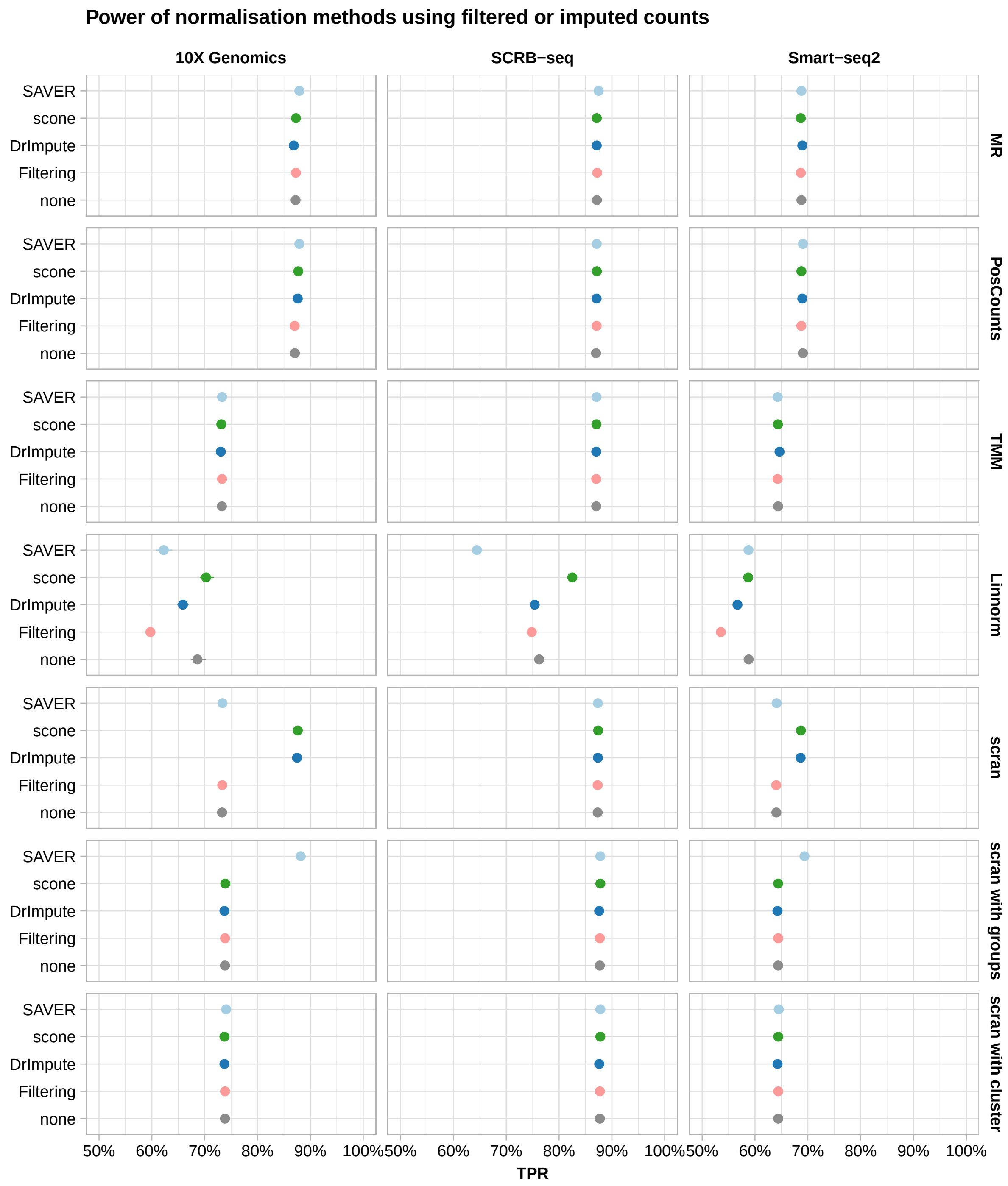


**Supplementary Figure 16:** Power of normalisation method using filtered or imputed counts. The expression of 10,000 genes over 768 cells (384 cells per group) were simulated and 20% of the genes were differentially expressed following an asymmetric narrow gamma distribution. The power to detect DE (TPR) based on DE-testing with limma-trend per count preprocessing approach stratified over library preparation protocol and normalisation method is plotted (mean TPR ± s.d.).

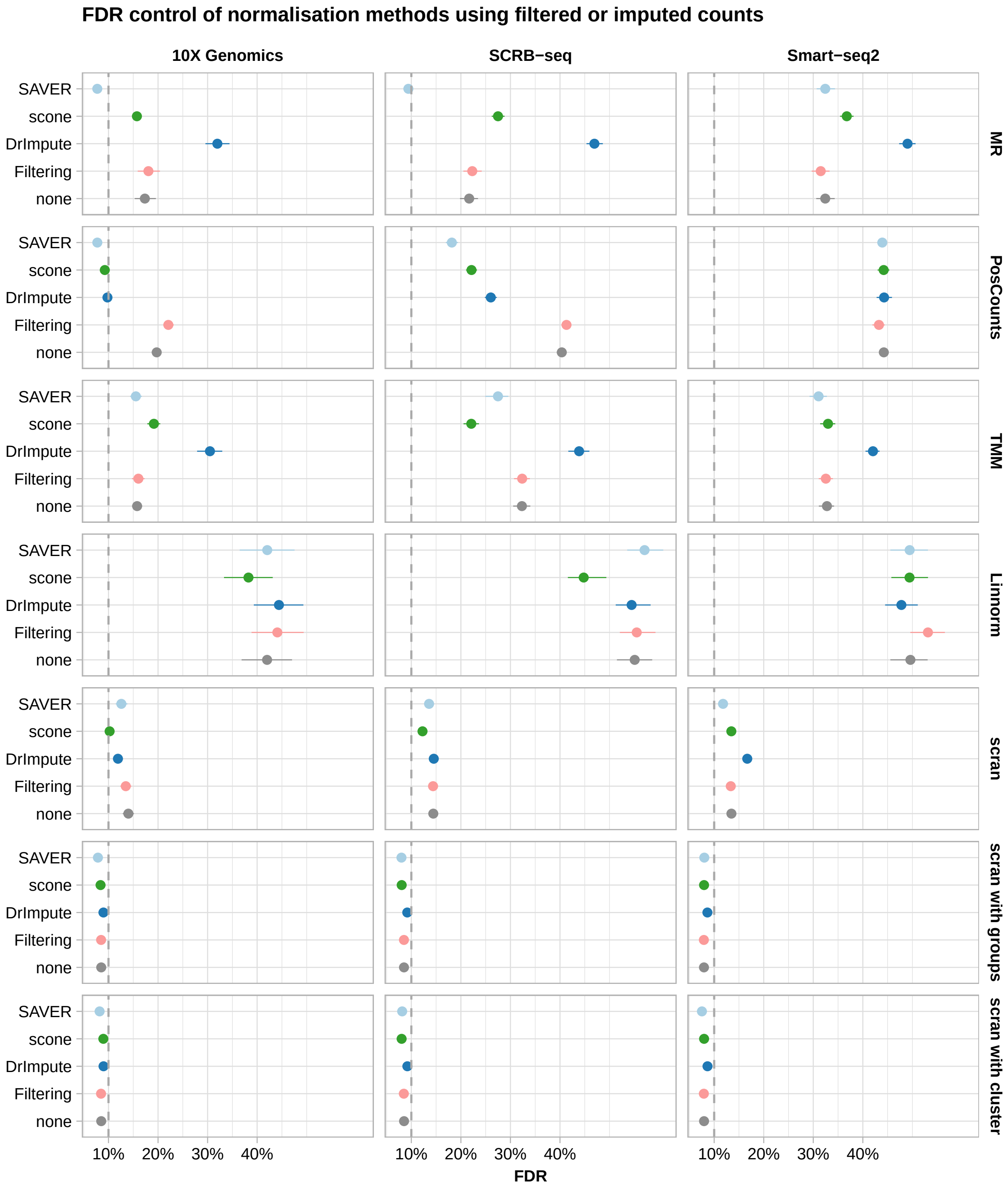


**Supplementary Figure 17:** FDR control of normalisation method using filtered or imputed counts. The expression of 10,000 genes over 768 cells (384 cells per group) were simulated and 20% of the genes were differentially expressed following an asymmetric narrow gamma distribution. The BH-FDR control based on DE-testing with limma-trend per count preprocessing approach stratified over library preparation protocol and normalisation method is plotted (mean FDR ± s.d.). The dashed line indicates the nominal FDR level of 10%.

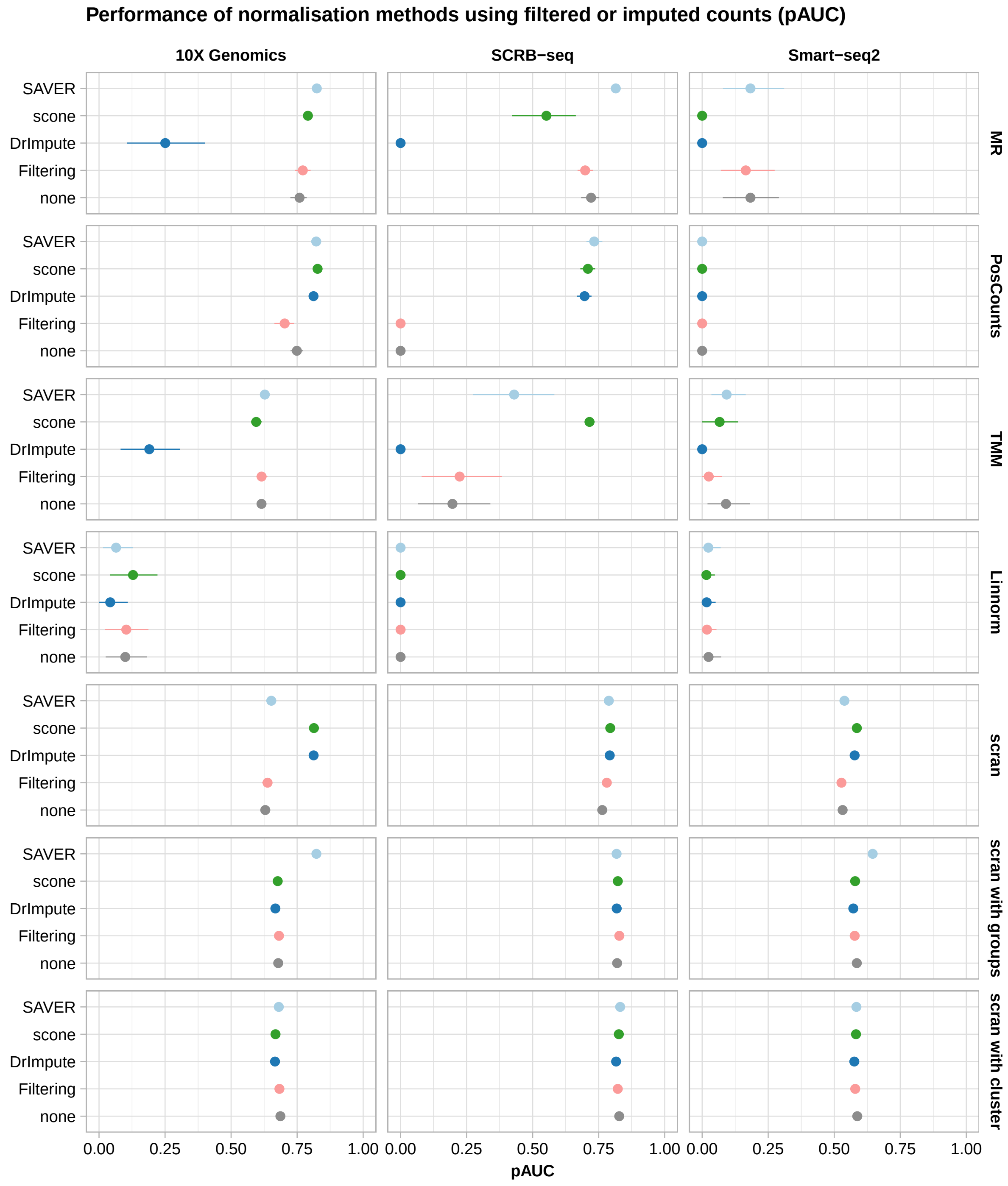


**Supplementary Figure 18:** Trade-off between power and false discoveries per normalisation method using filtered or imputed counts. The expression of 10,000 genes over 768 cells (384 cells per group) were simulated and 20% of the genes were differentially expressed following an asymmetric narrow gamma distribution. The discriminatory ability determined by the partial area under the curve (pAUC) based on DE-testing with limma-trend per count preprocessing approach stratified over library preparation protocol and normalisation method is plotted (mean pAUC $\pm$ s.d.).

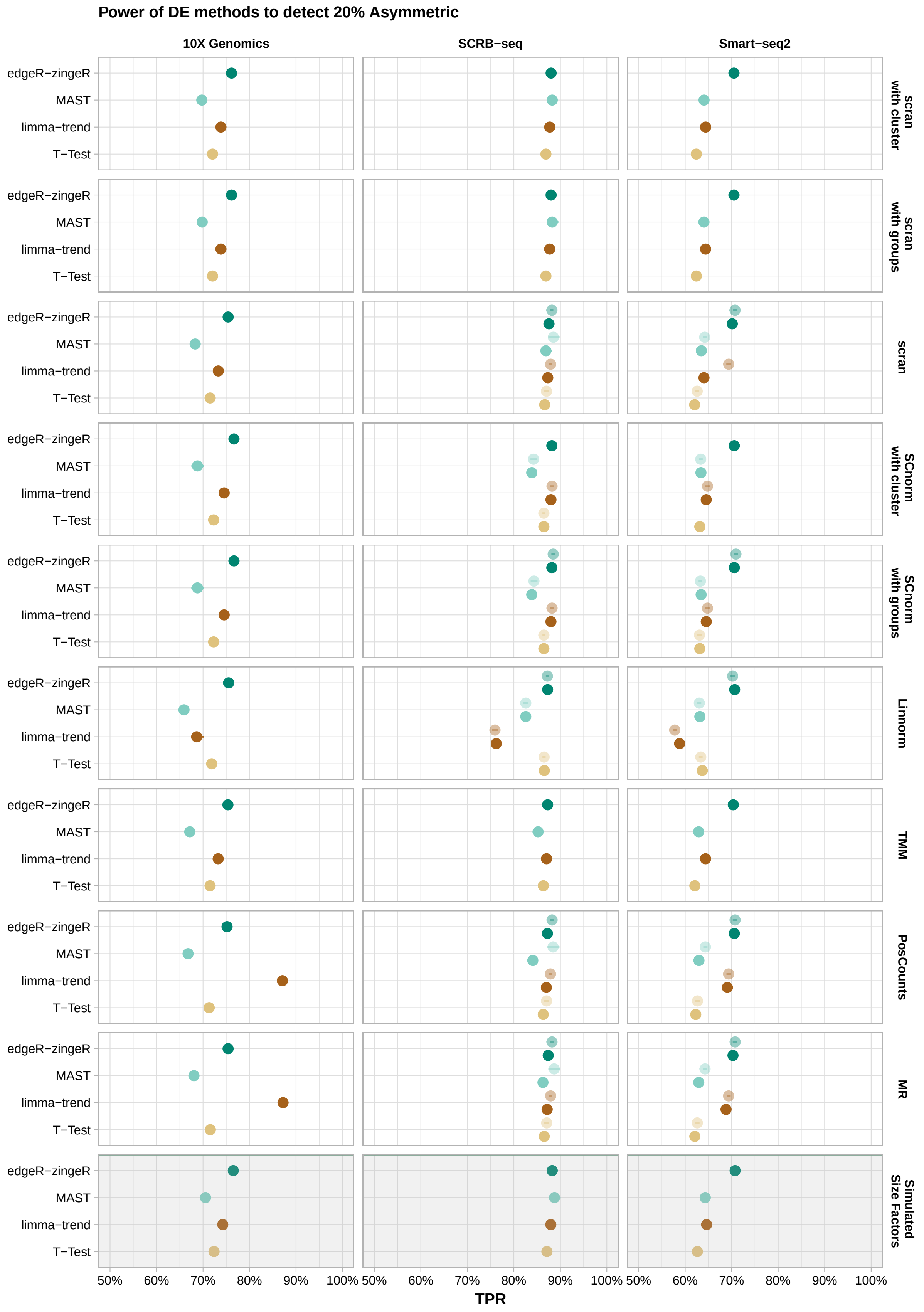


**Supplementary Figure 19:** Power of DE-tools for 20% Asymmetric. The expression of 10,000 genes over 768 cells (384 cells per group) were simulated given the observed mean-variance relation per protocol. 20% of the simulated genes are differentially expressed following an asymmetric narrow gamma distribution. Unfiltered counts were normalised using simulated library size factors or applying normalisation methods. Differential expression was tested using T-Test, limma-trend, MAST or edgeR-zingeR. The power to detect differential expression is plotted (mean TPR ± s.d.).

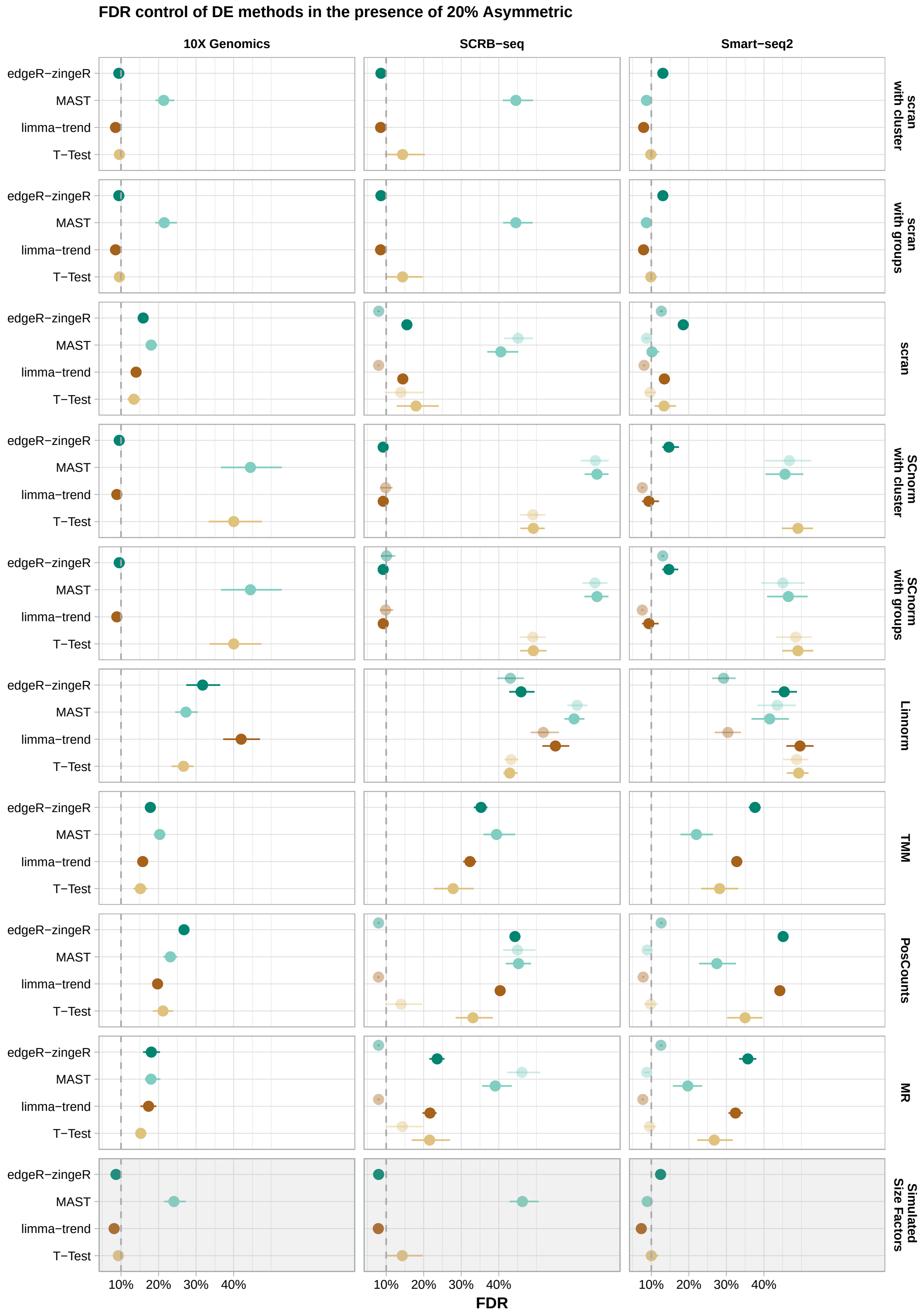


**Supplementary Figure 20:** FDR control of DE-tools for 20% Asymmetric. The expression of 10,000 genes over 768 cells (384 cells per group) were simulated given the observed mean-variance relation per protocol. 20% of the simulated genes are differentially expressed following an asymmetric narrow gamma distribution. Unfiltered counts were normalised using simulated library size factors or applying normalisation methods. Differential expression was tested using T-Test, limma-trend, MAST or edgeR-zingeR. The FDR control of the DE-methods is plotted (mean FDR ± s.d.). The dashed line indicates the nominal FDR level of 10%.

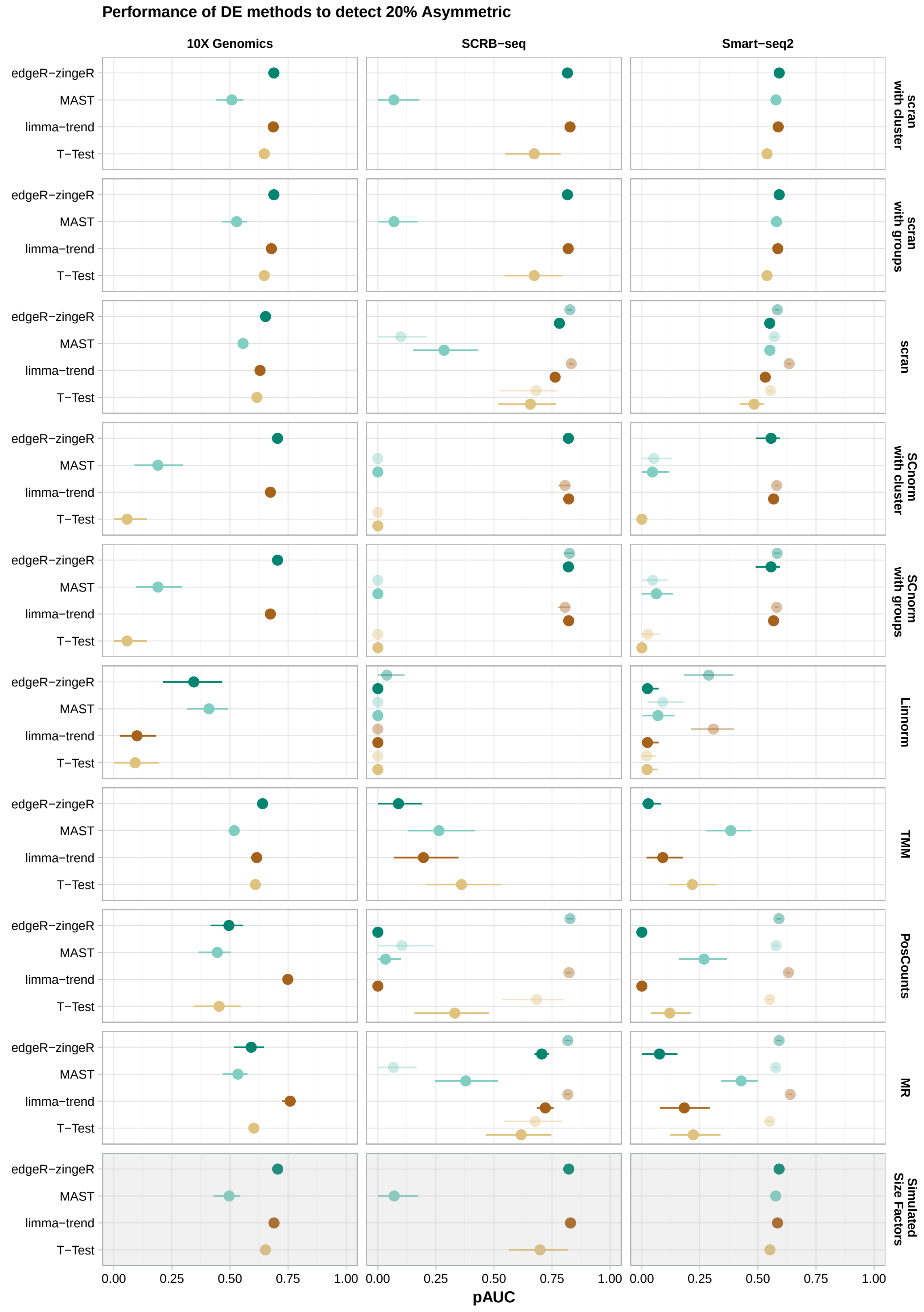


**Supplementary Figure 21:** The trade-off between power and false discoveries of DE-tools for 20% Asymmetric. The expression of 10,000 genes over 768 cells (384 cells per group) were simulated given the observed mean-variance relation per protocol. 20% of the simulated genes are differentially expressed following an asymmetric narrow gamma distribution. Unfiltered counts were normalised using simulated library size factors or applying normalisation methods. Differential expression was tested using T-Test, limma-trend, MAST or edgeR-zingeR. The discriminatory ability determined by the partial area under the curve (pAUC) based on the TPR-FDR curve is plotted (mean pAUC ± s.d.).

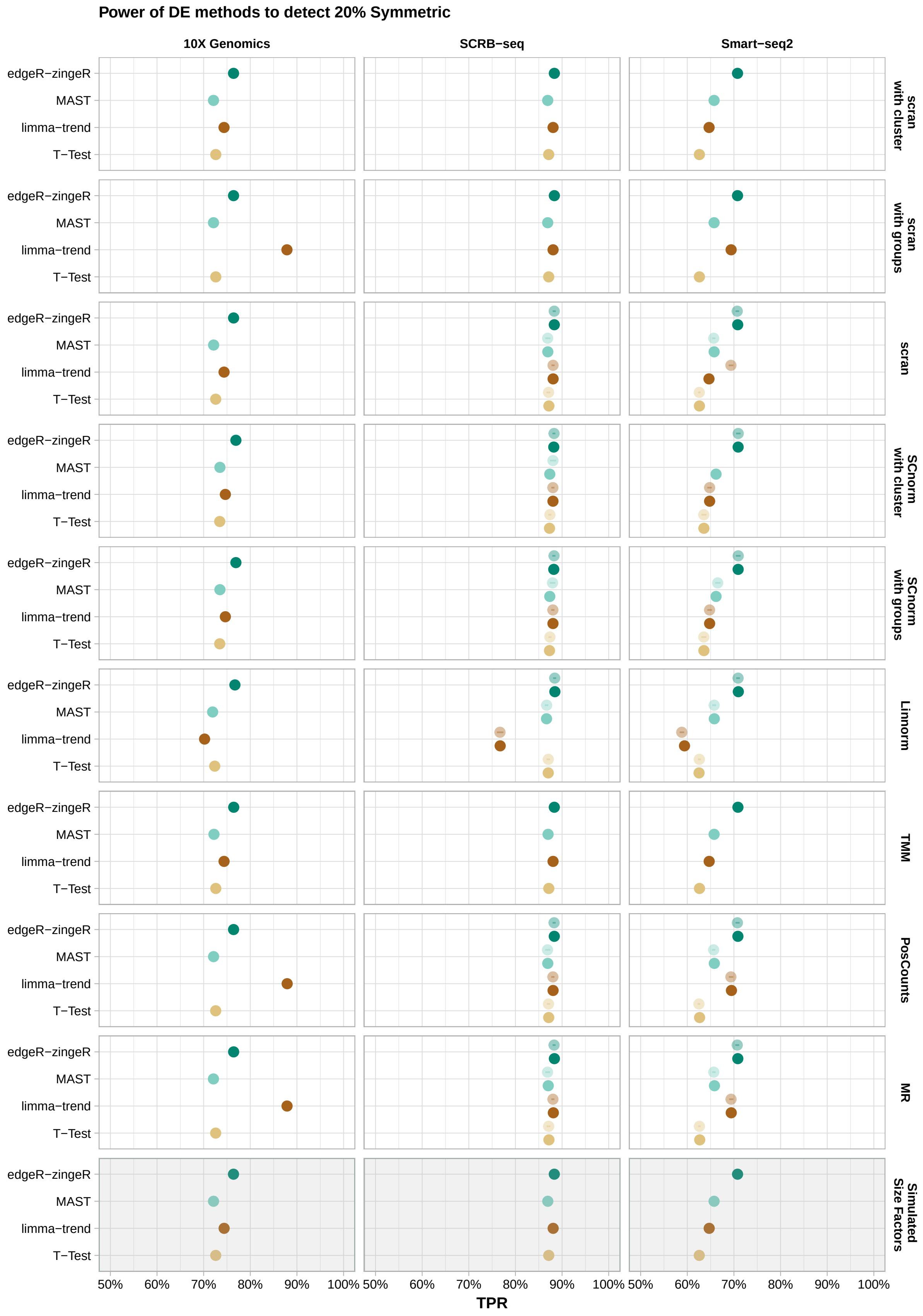


**Supplementary Figure 22:** Power of DE-tools for 20% Symmetric. The expression of 10,000 genes over 768 cells (384 cells per group) were simulated given the observed mean-variance relation per protocol. 20% of the simulated genes are differentially expressed following a symmetric narrow gamma distribution. Unfiltered counts were normalised using simulated library size factors or applying normalisation methods. Differential expression was tested using T-Test, limma-trend, MAST or edgeR-zingeR. The power to detect differential expression is plotted (mean TPR ± s.d.).

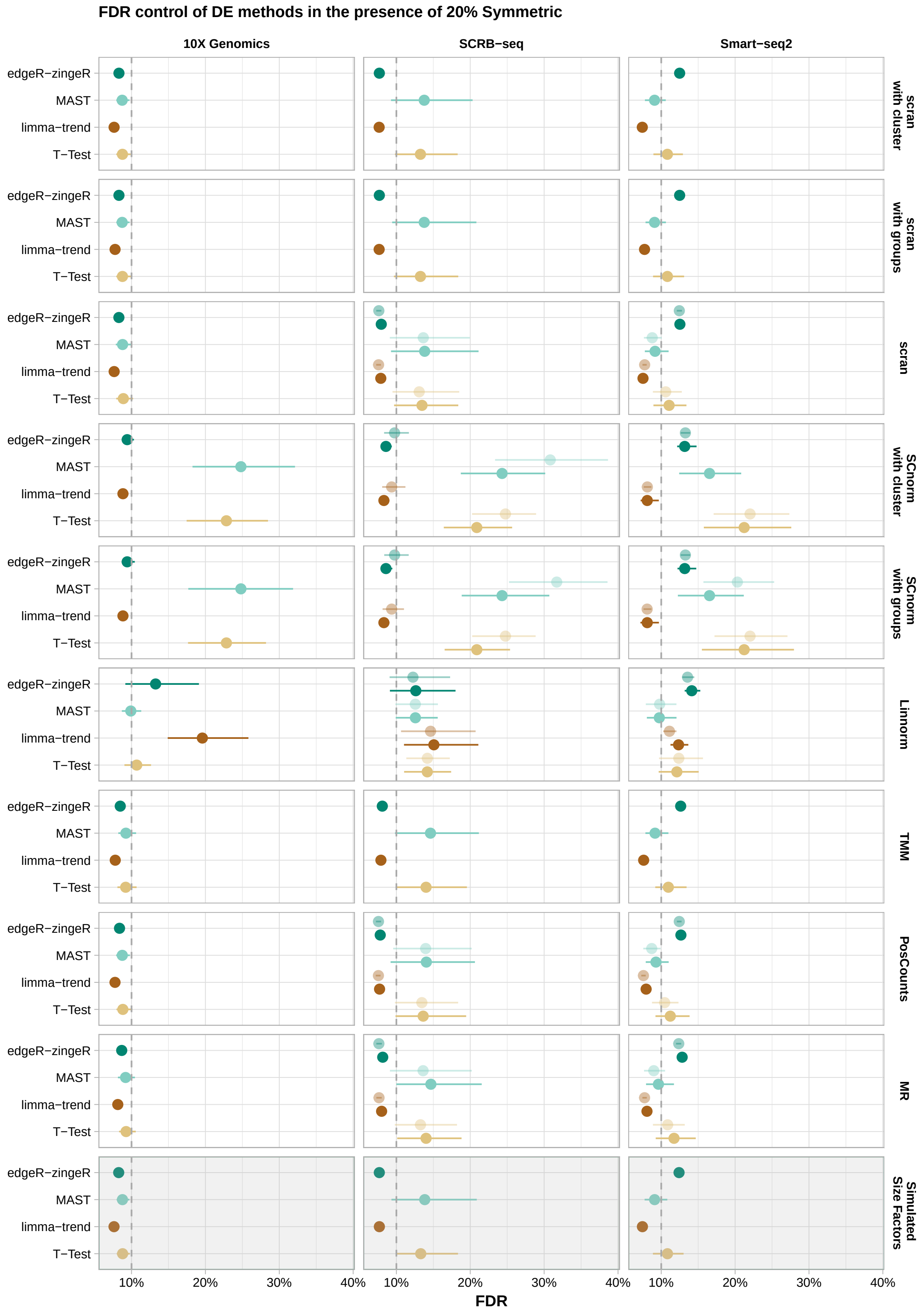


**Supplementary Figure 23:** FDR control of DE-tools for 20% Symmetric. The expression of 10,000 genes over 768 cells (384 cells per group) were simulated given the observed mean-variance relation per protocol. 20% of the simulated genes are differentially expressed following a symmetric narrow gamma distribution. Unfiltered counts were normalised using simulated library size factors or applying normalisation methods. Differential expression was tested using T-Test, limma-trend, MAST or edgeR-zingeR. The FDR control of the DE-methods is plotted (mean FDR ± s.d.). The dashed line indicates the nominal FDR level of 10%.

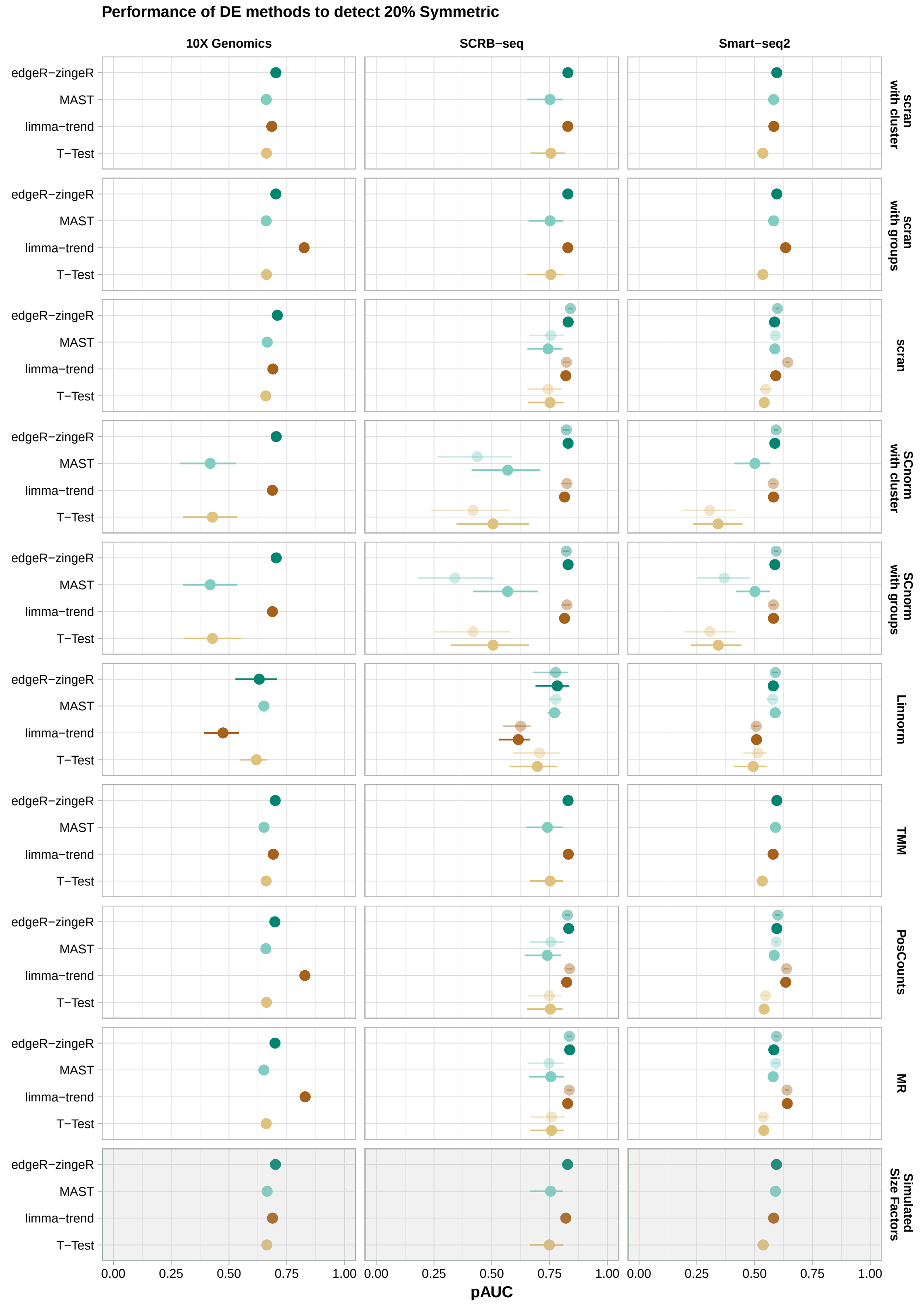


**Supplementary Figure 24:** The trade-off between power and false discoveries of DE-tools for 20% Symmetric. The expression of 10,000 genes over 768 cells (384 cells per group) were simulated given the observed mean-variance relation per protocol. 20% of the simulated genes are differentially expressed following a symmetric narrow gamma distribution. Unfiltered counts were normalised using simulated library size factors or applying normalisation methods. Differential expression was tested using T-Test, limma-trend, MAST or edgeR-zingeR. The discriminatory ability determined by the partial area under the curve (pAUC) based on the TPR-FDR curve is plotted ( mean pAUC ± s.d.).

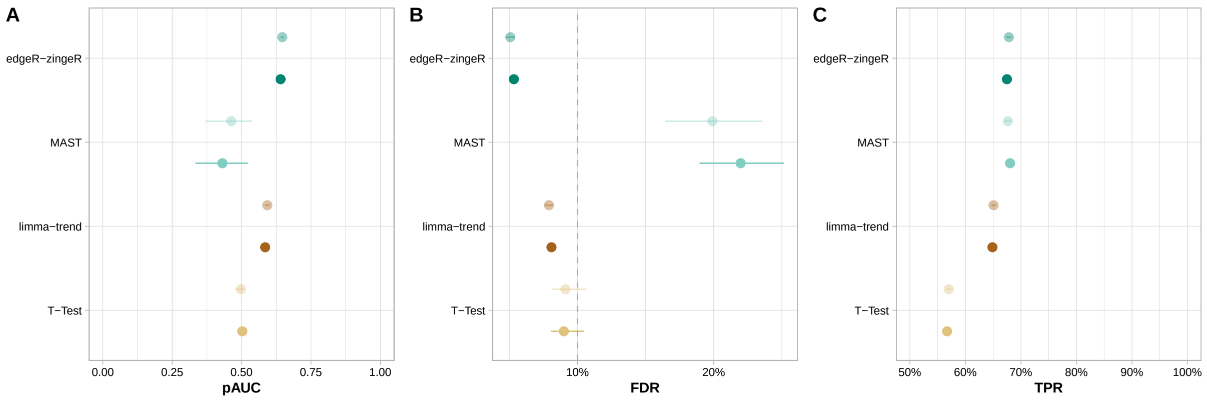


**Supplementary Figure 25:** Performance of DE-tools using Census normalisation. The expression of 10,000 genes over 768 cells (384 cells per group) were simulated given the observed mean-variance relation of Smart-seq2 data. 20% of the simulated genes are differentially expressed following an asymmetric narrow gamma distribution. Unfiltered counts were normalised using Census method. Differential expression was tested using T-Test, limma-trend, MAST or edgeR-zingeR. The lighter shade indicates the usage of spike-ins for normalisation.
A) The discriminatory ability determined by the partial area under the curve (pAUC) based on the TPR-FDR curve is plotted (mean pAUC ± s.d.). B) FDR control (mean FDR ± s.d.). The dashed line indicates the nominal FDR level of 10%. C) The power (TPR) to detect differential expression (mean TPR ± s.d.).

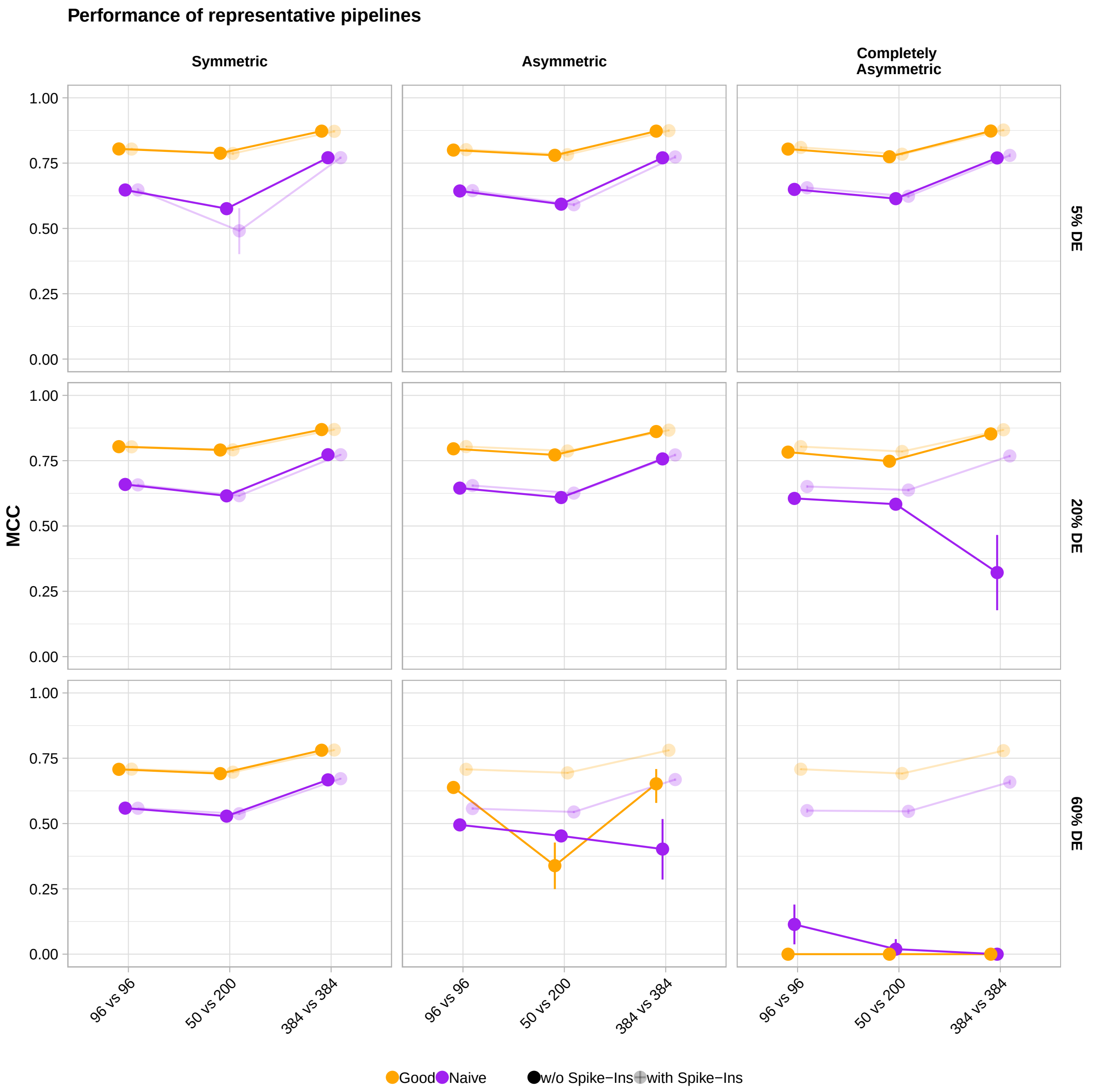


**Supplementary Figure 26:** Performance of pipelines. The expression of 10000 genes in 184, 250 or 768 cells were simulated and 5%, 20% or 60% of the genes were differentially expressed following a symmetric, asymmetric or completely asymmetric narrow gamma distribution. This simulation setup was applied to one good pipeline (SCRB-seq + STAR + GENCODE + no preprocessing + scran + limma-trend) and one naive pipeline (SCRB-seq + STAR + GENCODE + no preprocessing + MR + T-Test). For each analysis set, the Matthew Correlation Coefficient was averaged over 20 simulations (mean MCC ± s.d.) and rescaled to [0,1] interval.

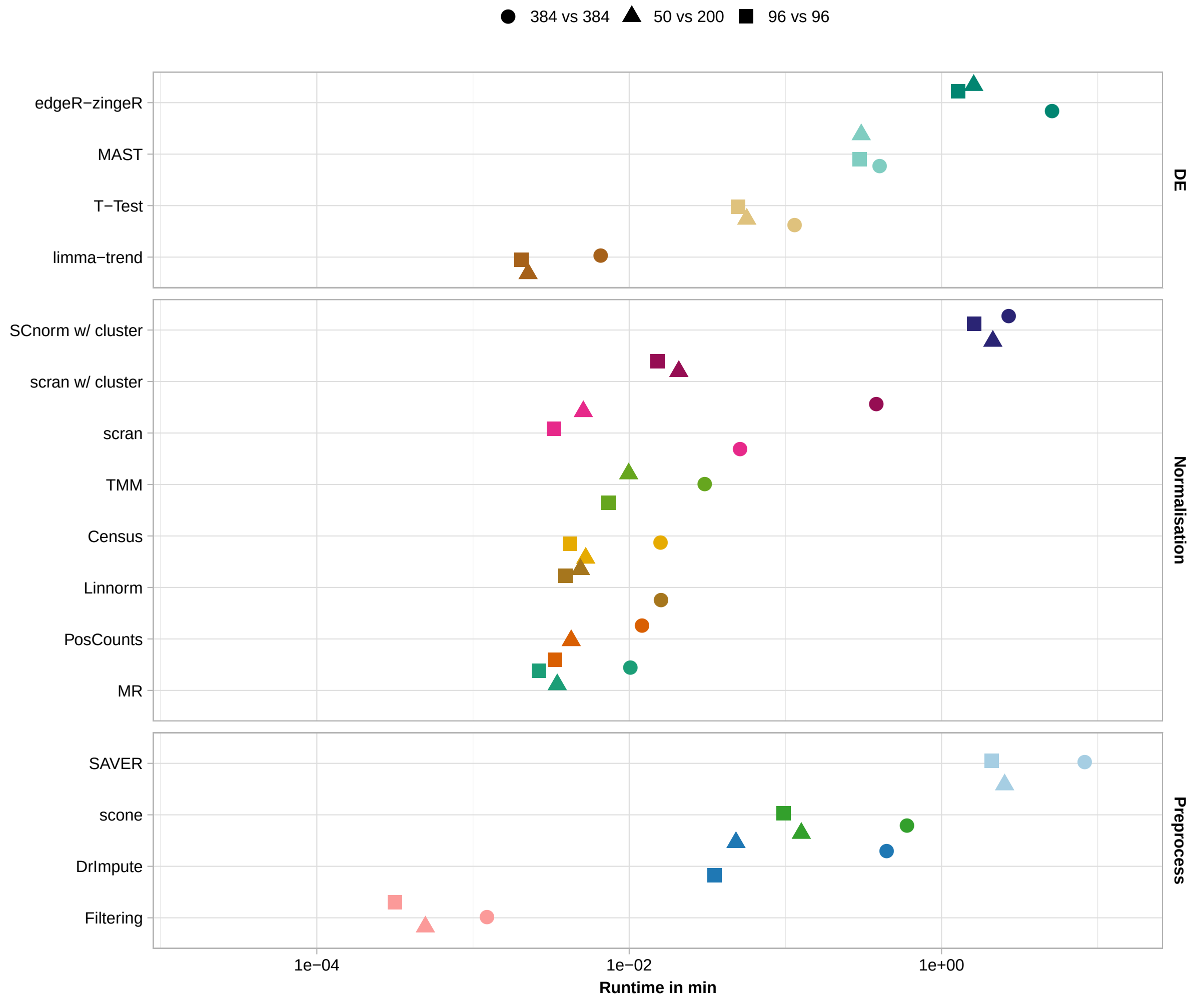


**Supplementary Figure 27:** Computational Run Time. The real CPU-time per pipeline step and method stratified over sample size.

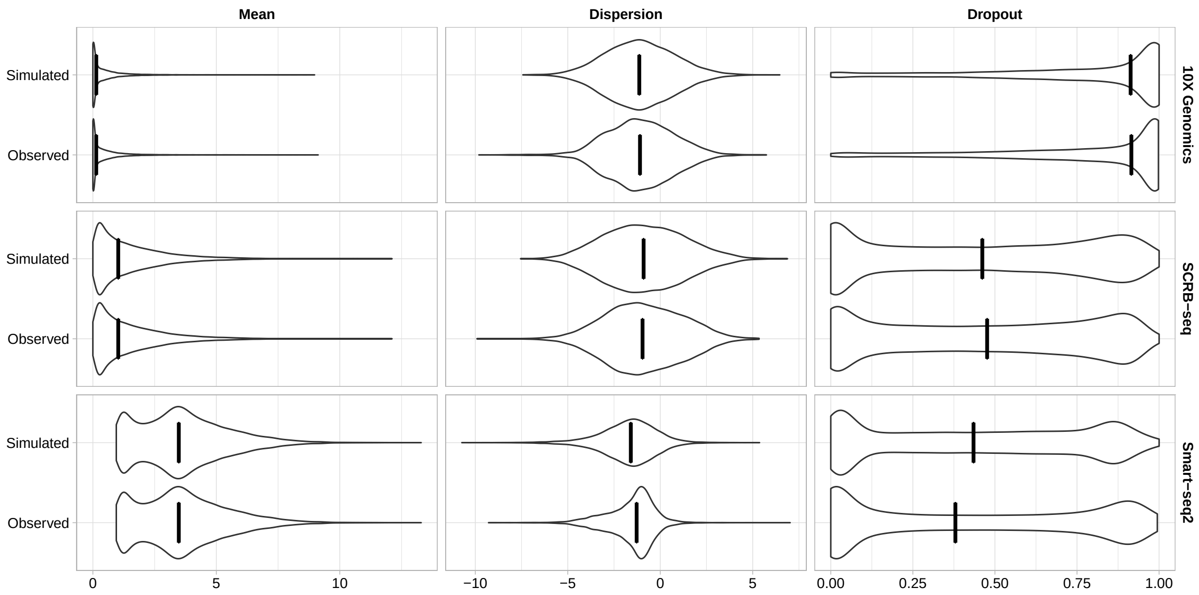


**Supplementary Figure 28:** Comparison of simulated and observed parameters. The marginal distribution of the observed and simulated log2 mean, log2 dispersion and gene dropout rate for Smart-seq2, SCRB-seq and 10X Genomics HGMM scRNA-seq data. For Smart-seq2 the mean and dispersion value excluding zeroes is plotted since a ZINB distribution is assumed. Black line indicates the median value.

### Supplementary Tables

| Protocol | Description | Cell Processing | Full Length | UMI | Number of cells | ERCC Spike-ins | Raw reads per cell |
|---|---|---|---|---|---|---|---|
| CEL-seq2 | J1 mESC cultured in 2i/LIF medium (two batches) | Fluidigm C1 | - | + | 48 + 48 | + | 1 million |
| Drop-seq | J1 mESC cultured in 2i/LIF medium (two batches) | Droplets | - | + | 45 + 34 | - | 1 million |
| SCRB-seq | J1 mESC cultured in 2i/LIF medium (two batches) | FACS | - | + | 44 + 49 | + | 1 million |
| Smart-seq2 | J1 mESC cultured in 2i/LIF medium (two batches) | FACS | + | - | 40 + 45 | + | 1 million |
| 10X Genomics | NH3T3 mouse cells (originally 1:1 mixture of human and mouse cells with a total of 1k cells) | 10X Genomics Chromium | - | + | 473 | - | ~60 thousand |
| 10X Genomics | Peripheral blood mononuclear cells (PBMC) | 10X Genomics Chromium | - | + | 1022 | - | ~54 thousand |

**Supplementary Table 1:** Description of single cell RNA-sequencing data sets.

| Name | Version | Number of Transcripts | Number of Genes | Number of Exons per Transcript | Number of Transcripts per Gene | Transcript Length | Gene Length |
|---|---|---|---|---|---|---|---|
| GENCODE | M15 | 131195 | 52636 | 6 | 2 | 1690 | 2348 |
| Vega | 68 | 110696 | 41175 | 6 | 3 | 1718 | 2671 |
| RefSeq (curated) | 85 | 34890 | - | 9 | - | 2852 | - |

**Supplementary Table 2:** Description of *Mus musculus* transcript annotations.

| Name | Command | Version | Ref. |
|---|---|---|---|
| BWA | bwa index -p [annotation_transcriptome_ercc.fa] | 0.7.12 | [9] |
| kallisto | kallisto index -i [annotation_transcriptome_ercc.fa] | 0.43.1 | [5] |
| BWA | bwa aln -t [bwa-index] [protocol_cDNA_reads.fastq] > [protocol.reads.sai] | 0.7.12 | [9] |
| BWA | bwa samse [reads.sai] [protocol_cDNA_reads.fastq] > [aligned.sam] | 0.7.12 | [9] |
| kallisto &Smart-seq2 | kallisto pseudo -i [kallisto-index] -o [outputpath] -b [protocol_cDNA_reads.fastq] --single -l [Mean_FragmentLength_Protocol] -d [SD_FragmentLength_Protocol] | 0.43.1 | [5] |
| kallisto & UMI | kallisto pseudo -i [kallisto-index] -o [outputpath] -b [protocol_cDNA_reads.fastq] --single --umi -l [Mean_FragmentLength_Protocol] -d [SD_FragmentLength_Protocol] | 0.43.1 | [5] |
| STAR zUMIs | STAR --runThreadN 12 --runMode genomeGenerate --genomeDir /index/star/[annotation] --genomeFastaFiles [mm10_genome_annotation_ercc.fa] --sjdbGTFfile [mm10_genome_annotation_ercc.gtf] --sjdbOverhang 44 | 2.5.3a | [10] |
| zUMIs | bash <path-to-zUMIs>/zUMIs-master.sh -y parameters.yaml ter.sh -f [protocol_barcode_reads.fq.gz] -r [protocol_cDNA_reads.fq.gz] -n [protocol-batch] -g [star-index] -a [mm10_genome_annotation_ercc.gtf] -c [cell-barcode-range] -m [umi-barcode-range] -l [read-length] -b [expected-cell-barcodes.txt] -o [outputpath] -d 1000000 -R no -S yes -s 0 -i [zUMIs-pipeline-path] | 0.0.3 | [11] |

**Supplementary Table 3:** Alignment and assignment commands for expression quantification.

| Pipeline Step | Method Name | Version | Reference |
|---|---|---|---|
| Preprocessing | Gene Dropout Filtering | - | - |
| Preprocessing | DrImpute | 1.0 | [12] |
| Preprocessing | scone | 1.6.1 | [13] |
| Preprocessing | SAVER | 1.1.1 | [14] |
| Normalisation | MR in DESeq2 | 1.22.2 | [15] |
| Normalisation | PosCounts in DESeq2 | 1.22.2 | [15] |
| Normalisation | TMM in edgeR | 3.24.3 | [16] |
| Normalisation | Census in monocle | 2.10.1 | [17] |
| Normalisation | Linnorm | 2.6.1 | [18] |
| Normalisation | SCnorm | 1.4.3 | [19] |
| Normalisation | scran | 1.10.1 | [20] |
| DE-tool | T-Test in stats | 3.5.3 | [21] |
| DE-tool | limma-trend | 3.38.3 | [22] |
| DE-tool | MAST | 1.8.2 | [23] |
| DE-tool | edgeR-zingeR | 0.1.0 | [24] |

**Supplementary Table 4:** Description of implemented methods in powsimR.
